# Global variation in prior exposure shapes antibody neutralization profiles of SARS-CoV-2 variants up to BA.2.86

**DOI:** 10.1101/2024.03.27.586820

**Authors:** Sam Turner, Gayatri Amirthalingam, Dalan Bailey, Dan H. Barouch, Kevin R. Bewley, Kevin Brown, Yunlong Cao, Yung-Wai Chan, Sue Charlton, DOVE consortium, Naomi S. Coombes, Bassam Hallis, David D. Ho, Fanchong Jian, Ninaad Lasrado, Ria Lassaunière, Lihong Liu, David C. Montefiori, Paul Moss, Joseph Newman, Helen Parry, Charlotta Polacek, Morten Rasmussen, Fei Shao, Xiaoying Shen, Nazia Thakur, Emma C. Thomson, Jing Wang, Peng Wang, Qian Wang, Brian J. Willett, Ayijiang Yisimayi, Derek J. Smith

**Affiliations:** Center for Pathogen Evolution, Department of Zoology, University of Cambridge, Cambridge, UK; UK Health Security Agency (UKHSA), London, UK; The Pirbright Institute, Woking, UK; Beth Israel Deaconess Medical Center, Boston, MA, USA; Medical Interventions Group, UK Health Security Agency, Porton Down, Salisbury SP4 0JG, UK; Biomedical Pioneering Innovation Center (BIOPIC), Peking University, Beijing, China; Changping Laboratory, Beijing, China; Aaron Diamond AIDS Research Center, Columbia University Vagelos College of Physicians and Surgeons, New York, NY, USA; Department of Medicine, Columbia University Vagelos College of Physicians and Surgeons, New York, NY, USA; Department of Microbiology and Immunology, Columbia University Vagelos College of Physicians and Surgeons, NY, USA; Department of Virus & Microbiological Special Diagnostics, Statens Serum Institut, Copenhagen 2300, Denmark; Duke University Medical Center, Durham, NC, USA; Institute of Immunology and Immunotherapy, University of Birmingham, Birmingham B15 2TT, UK; MRC University of Glasgow Centre for Virus Research, Glasgow G61 1QH, UK; Queen Elizabeth University Hospital, NHS Greater Glasgow and Clyde, Glasgow, UK

## Abstract

The highly mutated SARS-CoV-2 variant, BA.2.86, and its descendants are now the most frequently sequenced variants of SARS-CoV-2. We analyze antibody neutralization data from eight laboratories from the UK, USA, Denmark, and China, including two datasets assessing the effect of XBB.1.5 vaccines, to determine the effect of infection and vaccination history on neutralization of variants up to and including BA.2.86, and produce antibody landscapes to describe these neutralization profiles. We find evidence for lower levels of immune imprinting on pre-Omicron variants in sera collected from Denmark and China, which may be explained by lower levels of circulation of the ancestral variant in these countries, and the use of an inactivated virus vaccine in China.

## Introduction

A new sublineage of SARS-CoV-2, now designated BA.2.86, was first sequenced in July 2023 (World Health Organisation 2023). It has since been reported widely, with the global prevalence of BA.2.86 and its subvariants standing at ∼90% as of February 2024 (Figure 1A) - primarily composed of the JN.1, a subvariant of BA.2.86 containing the L455S spike substitution, which is now the most frequently sequenced variant of the virus (Shu and McCauley 2017; Khare et al. 2021). This resulted in BA.2.86 being designated as a Variant of Interest by the WHO in November 2023 (World Health Organisation 2023). BA.2.86 is characterized by a high number of mutations in key antigenic sites (Figure 1B), which may be driving its success. As such, there has been great interest in characterizing the antigenic phenotype of the variant, to determine the extent to which it evades current population immunity (Lasrado, Collier, Hachmann, et al. 2023; Wang, Guo, Liu, et al. 2023; Lassaunière et al. 2023; Yang et al. 2023; Coombes, Bewley, Le Duff, Alami-Rahmouni, et al. 2023; Khan et al. 2023; Chalkias et al. 2023; Wang, Guo, Bowen, et al. 2023; Willett et al. 2023).

**Figure 1:**
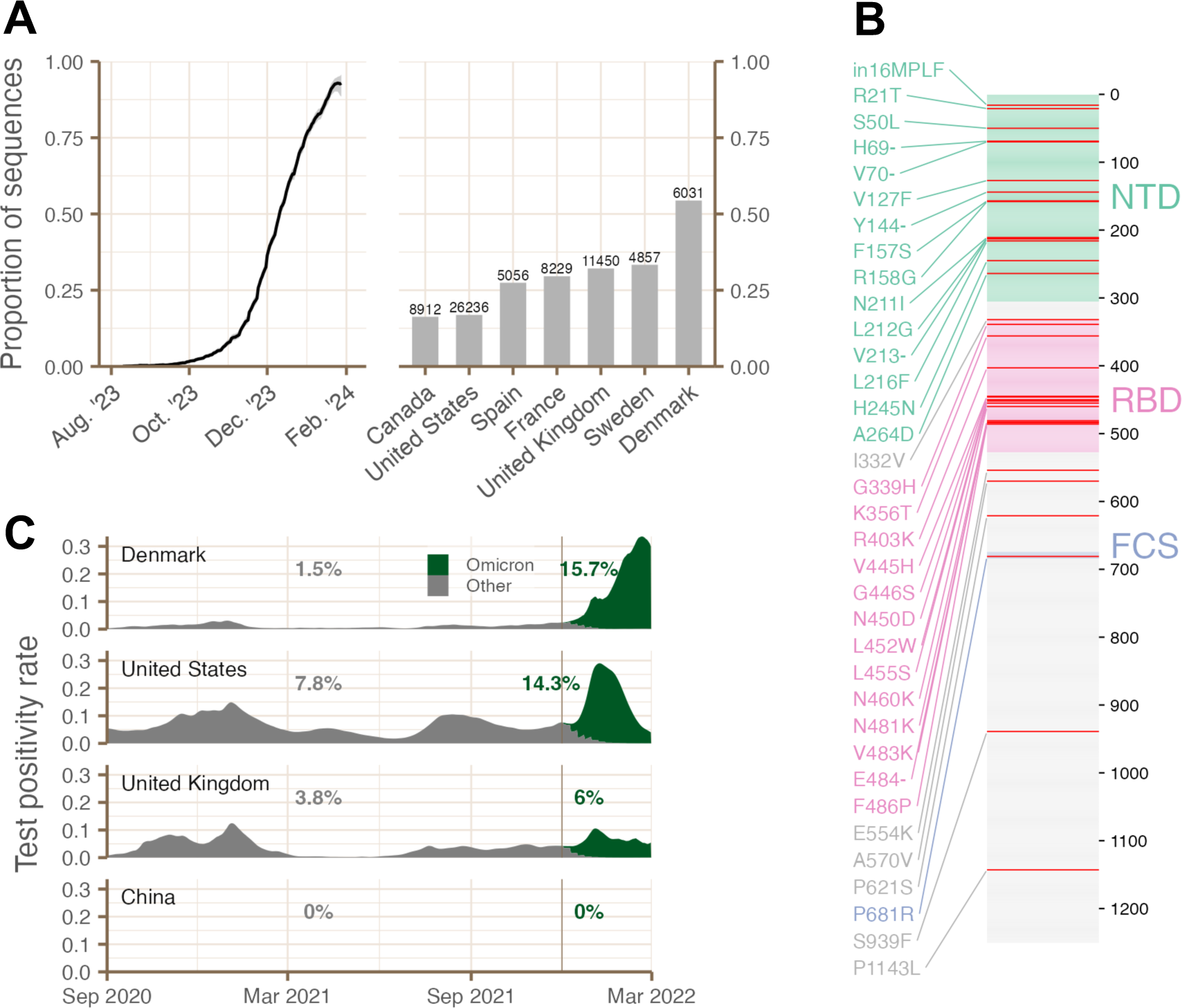
**(A)** Proportion of global sequences which are BA.2.86 (left), and proportion of sequences which are BA.2.86 in countries with >4000 BA.2.86 sequences submitted since 18 October 2023, with number of sequences written above each bar (right). **(B)** Mutations in the spike protein defining BA.2.86, relative to its ancestor BA.2. **(C)** Test positivity rate is shown for the period September 2020 to March 2022 (test positivity data is not available past this date), for each country where serum groups were taken from in this study, split into estimated Omicron and pre-Omicron positive tests. Numbers indicate the mean of the pre-Omicron test positivity rate (between 1^st^ September 2020 and 1^st^ December 2021) and Omicron test positivity rate (between 2^nd^ December 2021 and 1^st^ March 2022) in each country (the cutoff date of 1^st^ December 2021 is indicated with a vertical line). Details of the calculation of these quantities is given in the “Epidemiological data” section of the Methods. See Figure S4 for an analogous figure using number of confirmed infections.

Here we address this question by analyzing neutralization titers for serum cohorts from eight laboratories across the UK, USA, Denmark, and China, including two datasets assessing the effect of XBB.1.5 vaccines. We examine how differences in histories of prior variant exposure and vaccine uptake, which depend on the countries from which samples were taken, affect the neutralization profiles of individuals from these cohorts, including titers against BA.2.86.

## Results

Serum groups in this study are taken from several longitudinal studies, clinical cohorts, and one vaccine trial, containing a mixture of different histories of previous exposure to variants - predominantly BA.1/2/5 (“BA.x”) or XBB and subvariants of XBB (“XBB.x”) - both via infection and vaccination with bivalent BA.1 or BA.5 vaccines or with a monovalent XBB.1.5 vaccine. Further, neutralization titers were determined using different assay methodologies in the different laboratories, with a mixture of live virus neutralization and lentivirus-based or VSV-based pseudovirus neutralization assays, which limits direct comparability of data between studies (Table 1, Table S1). Geometric mean neutralization titers (GMT) for each serum group in this study are shown in Figure 2.

**Figure 2:**
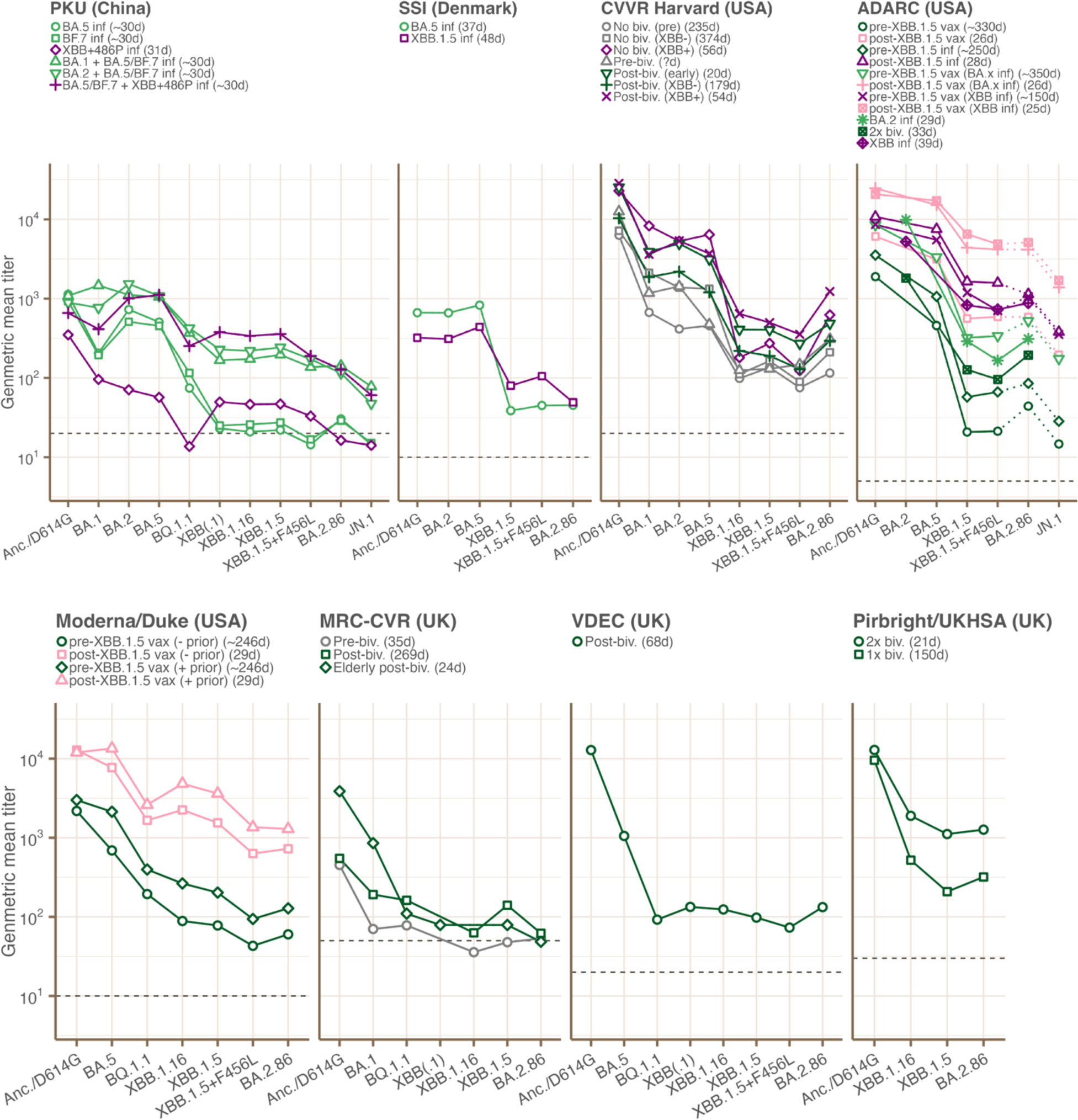
Geometric mean neutralizing antibody titers for each serum group included in the study, split by laboratory, with exposure history indicated by color. The number of days between the previous exposure (vaccination or infection) and the sample is given in parentheses in the legend for each serum group when known. In some serum groups from ADARC, titrations were not performed using BA.2.86, so titers were estimated from JN.1 titers for the purposes of producing antibody landscapes, as described in “Estimating BA.2.86 titers from JN.1 titers” in Methods. GMTs calculated from these estimated titers are included in this figure as small points connected with dotted lines. Dashed horizontal lines indicate the lower limit of detection of the assay used in each laboratory. Abbreviations are used in serum group names and exposure history: “vax” = vaccinated; “inf” = infected; “Anc.” = ancestral/Wuhan-1 SARS-CoV-2 virus; “biv.” = bivalent.

**Table 1:**
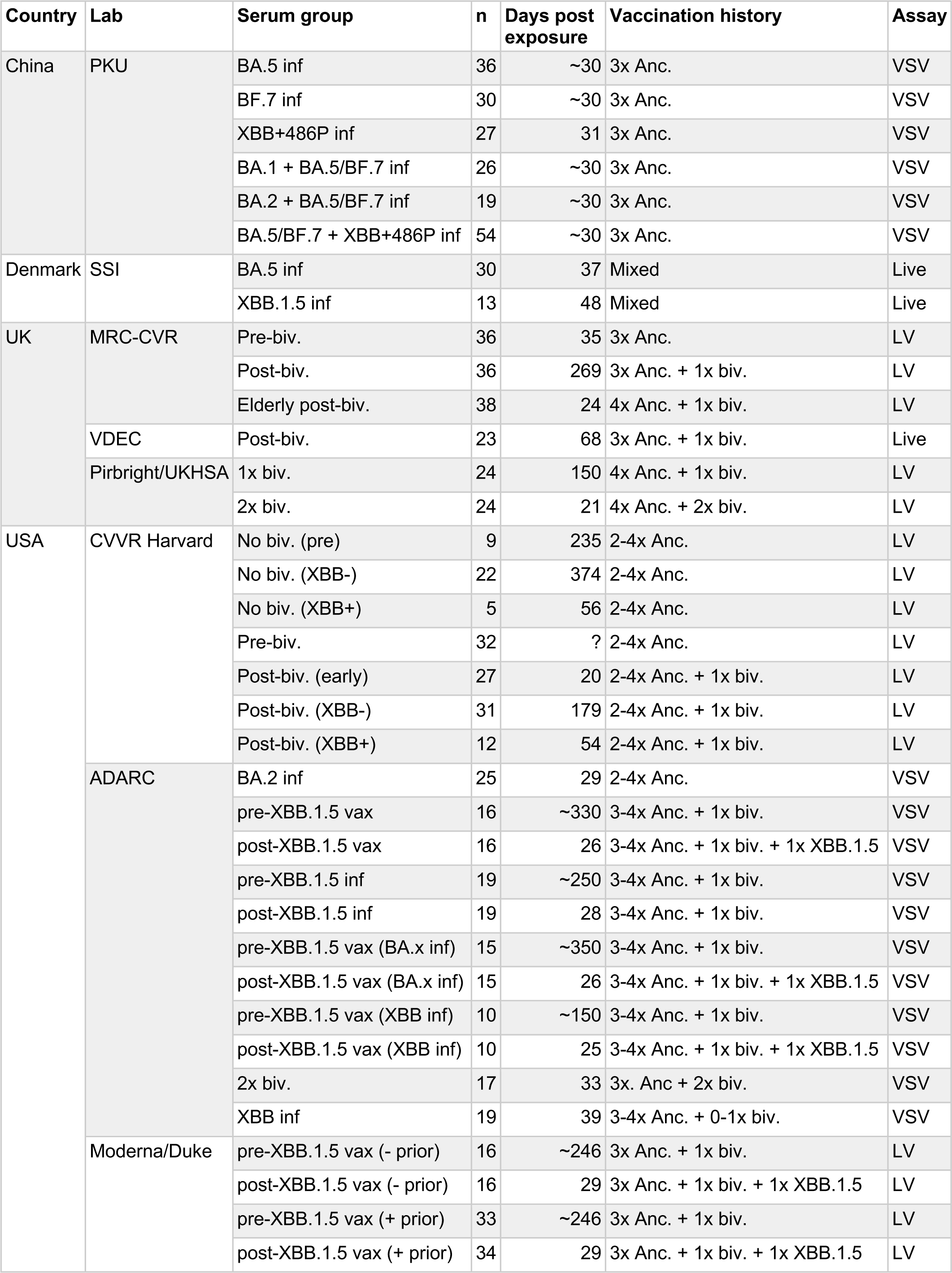
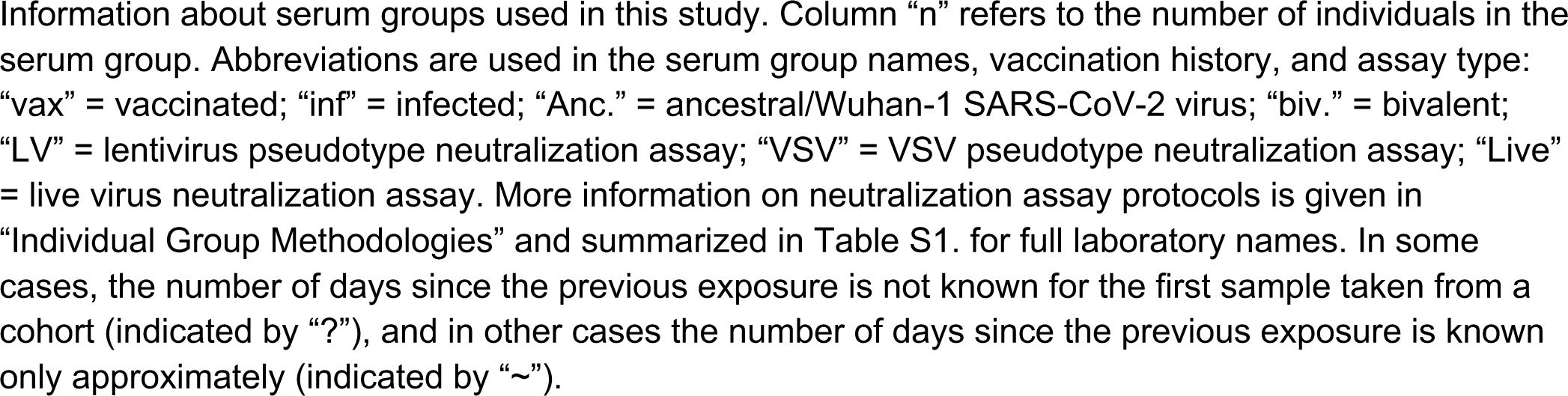

| Country | Lab | Serum group | n | Days post exposure | Vaccination history | Assay |
| --- | --- | --- | --- | --- | --- | --- |
| China | PKU | BA.5 inf | 36 | ~30 | 3x Anc. | VSV |
|  |  | BF.7 inf | 30 | ~30 | 3x Anc. | VSV |
|  |  | XBB+486P inf | 27 | 31 | 3x Anc. | VSV |
|  |  | BA.1 + BA.5/BF.7 inf | 26 | ~30 | 3x Anc. | VSV |
|  |  | BA.2 + BA.5/BF.7 inf | 19 | ~30 | 3x Anc. | VSV |
|  |  | BA.5/BF.7 + XBB+486P inf | 54 | ~30 | 3x Anc. | VSV |
| Denmark | SSI | BA.5 inf | 30 | 37 | Mixed | Live |
|  |  | XBB.1.5 inf | 13 | 48 | Mixed | Live |
| UK | MRC-CVR | Pre-biv. | 36 | 35 | 3x Anc. | LV |
|  |  | Post-biv. | 36 | 269 | 3x Anc. + 1x biv. | LV |
|  |  | Elderly post-biv. | 38 | 24 | 4x Anc. + 1x biv. | LV |
|  | VDEC | Post-biv. | 23 | 68 | 3x Anc. + 1x biv. | Live |
|  | Pirbright/UKHSA | 1x biv. | 24 | 150 | 4x Anc. + 1x biv. | LV |
|  |  | 2x biv. | 24 | 21 | 4x Anc. + 2x biv. | LV |
| USA | CVVR Harvard | No biv. (pre) | 9 | 235 | 2-4x Anc. | LV |
|  |  | No biv. (XBB-) | 22 | 374 | 2-4x Anc. | LV |
|  |  | No biv. (XBB+) | 5 | 56 | 2-4x Anc. | LV |
|  |  | Pre-biv. | 32 | ? | 2-4x Anc. | LV |
|  |  | Post-biv. (early) | 27 | 20 | 2-4x Anc. + 1x biv. | LV |
|  |  | Post-biv. (XBB-) | 31 | 179 | 2-4x Anc. + 1x biv. | LV |
|  |  | Post-biv. (XBB+) | 12 | 54 | 2-4x Anc. + 1x biv. | LV |
|  | ADARC | BA.2 inf | 25 | 29 | 2-4x Anc. | VSV |
|  |  | pre-XBB.1.5 vax | 16 | ~330 | 3-4x Anc. + 1x biv. | VSV |
|  |  | post-XBB.1.5 vax | 16 | 26 | 3-4x Anc. + 1x biv. + 1x XBB.1.5 | VSV |
|  |  | pre-XBB.1.5 inf | 19 | ~250 | 3-4x Anc. + 1x biv. | VSV |
|  |  | post-XBB.1.5 inf | 19 | 28 | 3-4x Anc. + 1x biv. | VSV |
|  |  | pre-XBB.1.5 vax (BA.x inf) | 15 | ~350 | 3-4x Anc. + 1x biv. | VSV |
|  |  | post-XBB.1.5 vax (BA.x inf) | 15 | 26 | 3-4x Anc. + 1x biv. + 1x XBB.1.5 | VSV |
|  |  | pre-XBB.1.5 vax (XBB inf) | 10 | ~150 | 3-4x Anc. + 1x biv. | VSV |
|  |  | post-XBB.1.5 vax (XBB inf) | 10 | 25 | 3-4x Anc. + 1x biv. + 1x XBB.1.5 | VSV |
|  |  | 2x biv. | 17 | 33 | 3x. Anc + 2x biv. | VSV |
|  |  | XBB inf | 19 | 39 | 3-4x Anc. + 0-1x biv. | VSV |
|  | Moderna/Duke | pre-XBB.1.5 vax (- prior) | 16 | ~246 | 3x Anc. + 1x biv. | LV |
|  |  | post-XBB.1.5 vax (- prior) | 16 | 29 | 3x Anc. + 1x biv. + 1x XBB.1.5 | LV |
|  |  | pre-XBB.1.5 vax (+ prior) | 33 | ~246 | 3x Anc. + 1x biv. | LV |
|  |  | post-XBB.1.5 vax (+ prior) | 34 | 29 | 3x Anc. + 1x biv. + 1x XBB.1.5 | LV |

### To what extent does BA.2.86 escape neutralization relative to previous variants?

To get a broad picture of the extent to which BA.2.86 escapes existing population immunity, we looked at the fold change in neutralization titer to BA.2.86 relative to a set of key historical variants (Figure 3). Neutralization titers to BA.2.86 are similar to those to XBB.1.5, with titers to XBB.1.5 an average of 1.2x higher than those to BA.2.86 across all serum groups (Figure 3A). However, this fold change differs between serum groups with different exposure histories (Figure 3B): titers to BA.2.86 are either not significantly different from or slightly higher than titers to XBB.1.5 for serum groups from the UK or USA without an exposure (via infection or vaccination) to XBB.x, for serum groups from China with one BA.x infection, or for serum groups from Denmark with a BA.5 infection. By contrast, titers to XBB.1.5 are significantly higher than titers to BA.2.86 for serum groups from the UK or USA with an XBB.x exposure, from China with two BA.x infections or an XBB.x infection, or from Denmark with an XBB.1.5 infection (fold reductions of 1.4x, 2.3x, 1.6x respectively) (Figure 3B).

**Figure 3:**
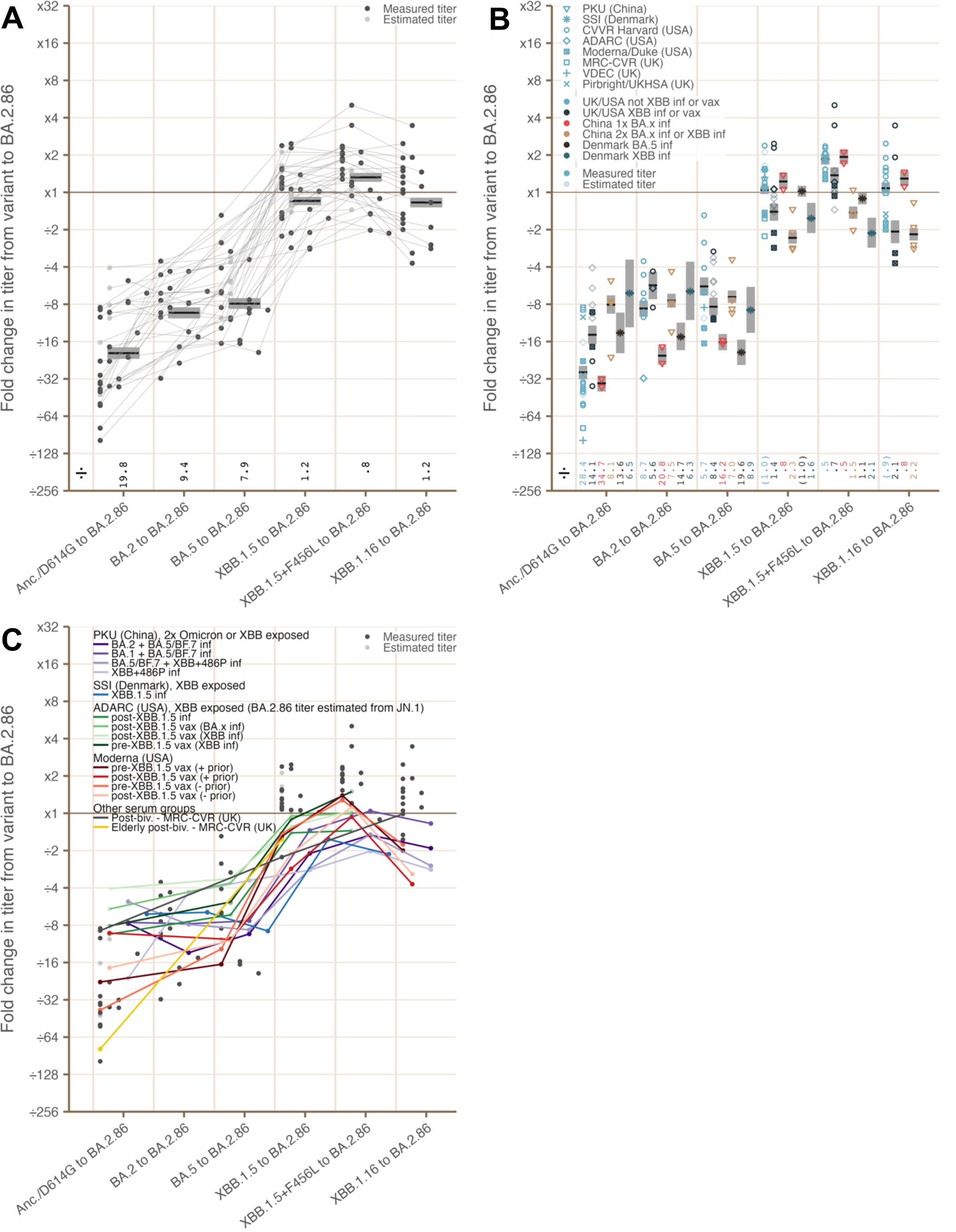
**(A)** Geometric mean fold change in neutralization titers against historic variants relative to BA.2.86. Individual points show mean fold change values for individual serum groups, and the black horizontal line shows the geometric mean fold change across all serum groups against a particular variant (also written along the x-axis), and the gray shading indicates a 95% confidence interval for the mean calculated by the non-parametric bootstrap using 1000 replicates. Points calculated using estimated BA.2.86 titers are shown in lighter color (see “Estimating BA.2.86 titers from JN.1 titers” in Methods) **(B)** Same as **A**, with mean values shown for different locations and exposure histories. Fold changes which are not significant at the 5% level by the Wilcoxon paired difference test are shown in parentheses. **(C)** Same as **A**, with serum groups showing high fold reduction in neutralization titer to BA.2.86 relative to XBB.1.5/16 highlighted (see Methods for details of how these serum groups are chosen). Abbreviations are used in serum group names and exposure history: “vax” = vaccinated; “inf” = infected; “Anc.” = ancestral/Wuhan-1 SARS-CoV-2 virus; “biv.” = “bivalent.

Relative to pre-XBB variants, fold reduction across all serum groups from BA.5 to BA.2.86 was 7.9x, and from ancestral virus or D614G (Anc./D614G) to BA.2.86 was 19.8x (Figure 3A). Again, fold reduction in titer from Anc./D614G to BA.2.86 varied according to exposure history: serum groups without a confirmed XBB.x exposure in the UK or USA had a fold reduction of 28.4x, compared to 6.5-14.1x for UK or USA serum groups with an XBB.x exposure, serum groups from Denmark with an XBB.1.5 or BA.5 infection, and serum groups from China with two BA.x infections or an XBB.x infection (Figure 3B).

### Which serum groups have lower neutralization titers to BA.2.86 than to XBB.1.5 and XBB.1.16?

To better understand the variation in BA.2.86 neutralization titers, we identified serum groups that neutralize BA.2.86 less effectively relative to XBB.1.5 or XBB.1.16 (Figure 3C). The majority of these serum groups fall into four categories which are discussed further below: (1) serum groups from China with two BA.x infections or with an XBB.x infection; (2) the serum group from Denmark with an XBB.1.5 infection; (3) serum groups from the ADARC (USA) XBB.1.5 vaccine cohorts which either have been infected or vaccinated with XBB.1.5 (note that, for ADARC (USA) serum groups, titers to BA.2.86 are estimated as 3x the titer to JN.1 for this comparison, as titrations were not performed to BA.2.86 - see Figure 2, and “Estimating BA.2.86 titers from JN.1 titers’’ in Methods for more discussion); and (4) serum groups from the Moderna XBB.1.5 vaccine trial dataset.

For these serum groups, we compared the fold reduction in titer from other variants to XBB.1.5, and found that it is typically smaller than in other serum groups, which suggests that XBB.1.5 titers are higher than typical in these serum groups (Figure 4A). We also compared fold reduction from non-XBB variants to BA.2.86 for these serum groups, and found that it is not larger than average, and is lower than average relative to D614G for most of these serum groups - which further suggests that these serum groups have higher than typical XBB.1.5/16 titers, rather than lower than typical BA.2.86 titers (potential reasons for which are discussed later). Indeed, it should be noted that the absolute BA.2.86 titers of these groups are often higher than those of other groups measured in the same assay (Figure 2) - as demonstrated in the clinical cohort (ADARC) and Moderna XBB.1.5 vaccine trial where BA.2.86 titers increase substantially upon XBB.1.5 vaccination.

**Figure 4:**
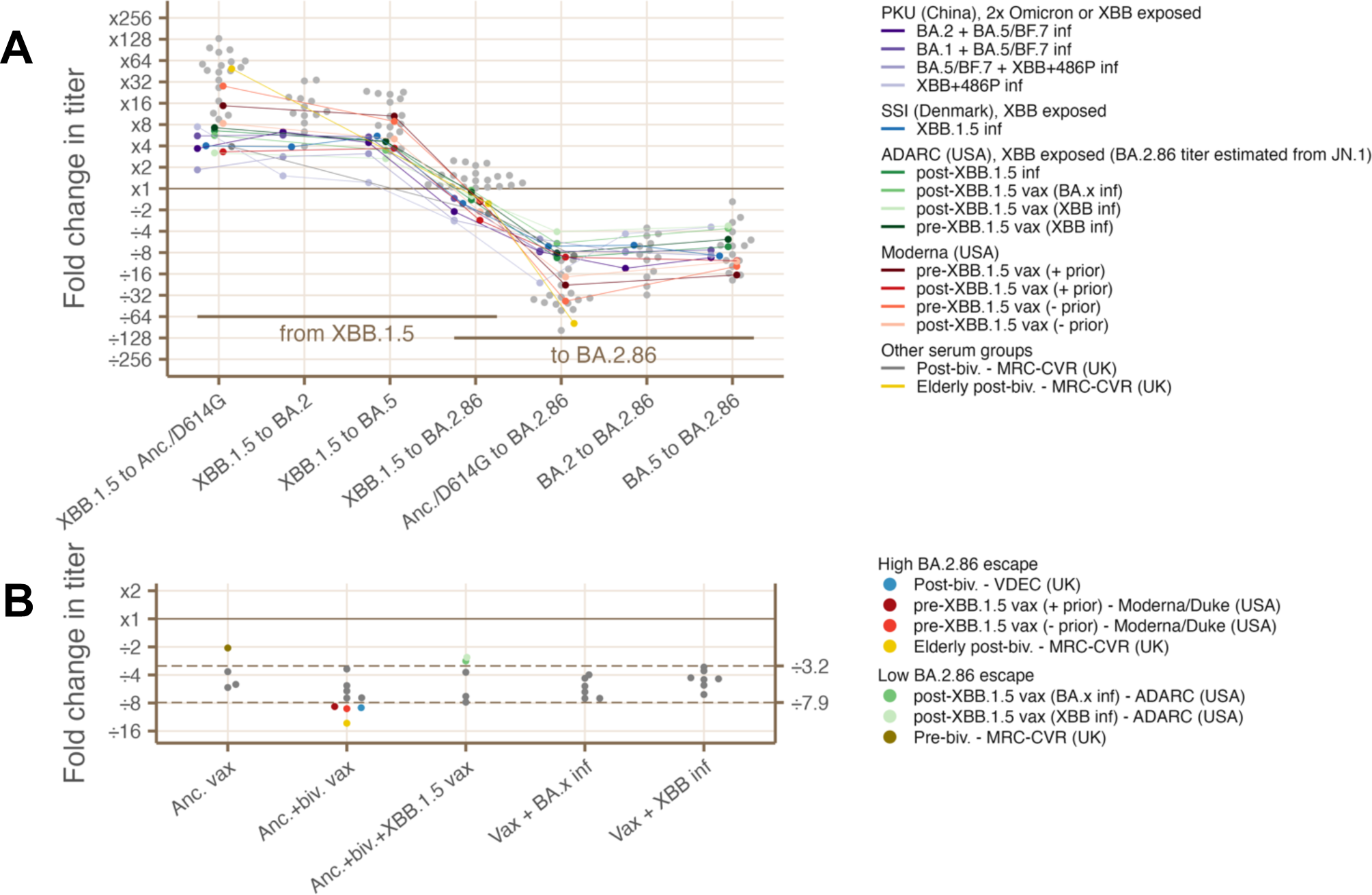
**(A)** Comparisons of fold change in neutralization titer for serum groups which show reductions in neutralization titer against BA.2.86 relative to XBB.1.5/16 (see Methods for details of how these serum groups are identified). Fold change values are shown from XBB.1.5 to each other variant, and then from each other variant to BA.2.86. Serum groups showing a reduction titer from BA.2.86 to XBB.1.5/16 are plotted in color, compared to all other serum groups which are plotted in gray. **(B)** Fold change from a mean across all pre-BA.2.86 variants to BA.2.86 (see Methods for details of this calculation), with serum groups showing a fold reduction outside of the range 3.2-7.9x plotted in color, compared to all other serum groups plotted as gray points. Abbreviations are used in serum group names: “vax” = vaccinated; “inf” = infected; “biv.” = bivalent.

### The peak of the antibody landscape is shifted for serum groups from China with two BA.x or BA.x plus XBB.x infections or from Denmark with one BA.5 or XBB.1.5 infection

In the three serum groups from China with two BA.x or BA.x plus XBB.x infections and the two serum groups from Denmark with an XBB.1.5 or BA.5 infection, there is little or no reduction in titers between Anc./D614G and BA.x variants (Figure 5C). This is reflected in the peak of the antibody landscape being shifted away from the Anc./D614G region of antigenic space and towards the BA.x region of antigenic space (Figures 5A, 5B). This shift in the peak of the landscape is not seen in serum groups from either the UK or USA, where the peak of the antibody landscape remains nearer to the original Anc./D614G variant regardless of infection or vaccination history.

**Figure 5:**
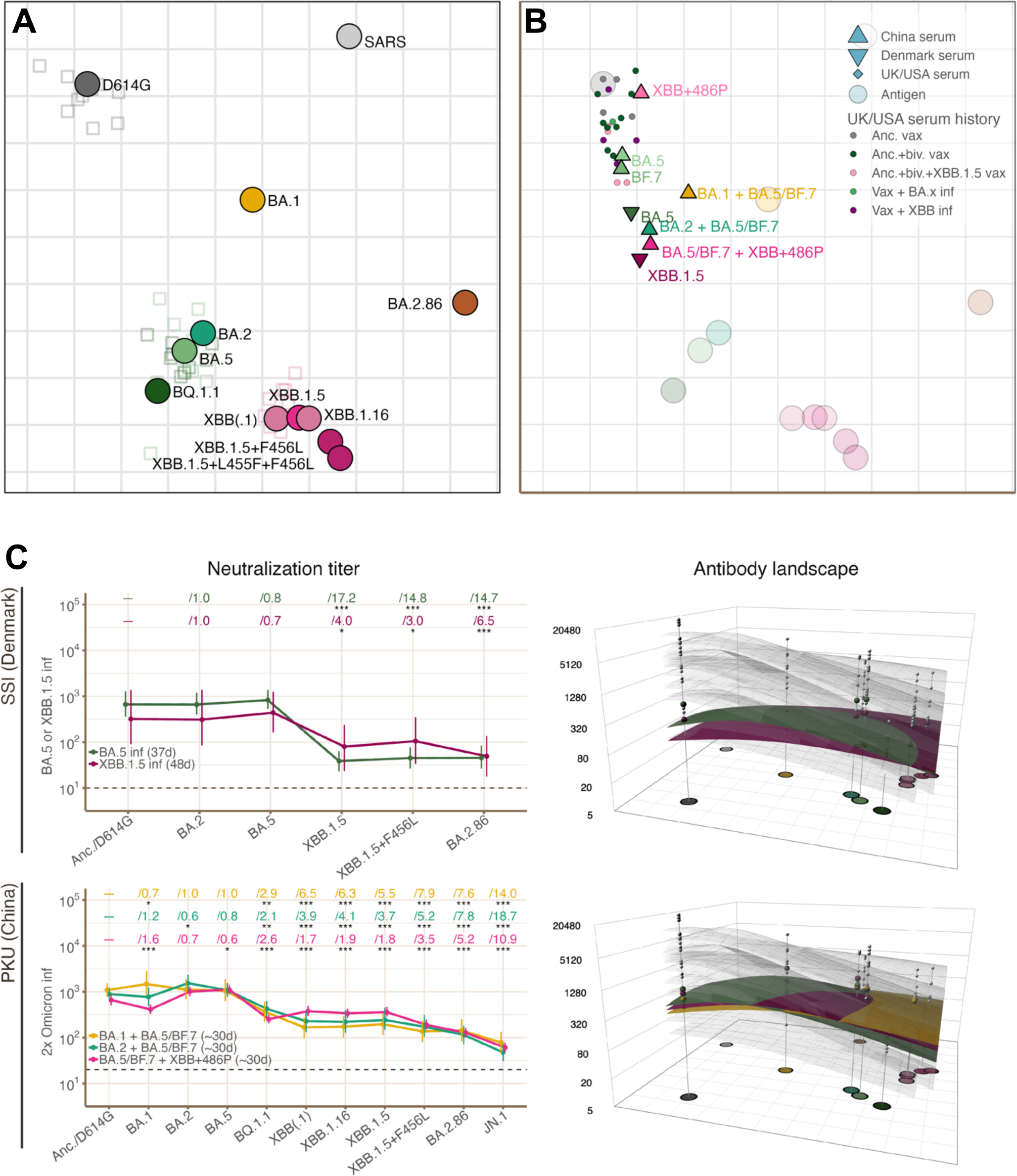
**(A)** Antigenic map used to construct antibody landscapes, produced from neutralization titration data for mouse antisera. **(B)** Location of the peaks of the antibody landscapes for all serum groups, with serum groups from Denmark and China highlighted. **(C)** Neutralization titers (left column) and antibody landscapes (right column) for serum groups with shifted peaks of reactivity from Denmark and China (colored landscapes), compared to BA.x infection serum groups from the UK and USA (pale gray landscapes). Fold reduction in titer relative to Anc./D614G is given for each variant in the left column, and asterisks indicate level of statistical significance (* = p<0.05; ** = p<0.01; *** = p<0.001, as determined by the Wilcoxon paired difference test). 95% confidence intervals are shown, calculated by the non-parametric bootstrap using 1000 replicates. The number of days between the previous exposure (vaccination or infection) and the sample is given in parentheses in the legend for each serum group when known. Abbreviations are used in serum group names and exposure history: “vax” = vaccinated; “inf” = infected; “Anc.” = ancestral/Wuhan-1 SARS-CoV-2 virus; “biv.” = bivalent.

The shift in the peak of reactivity towards the infecting variant for these serum groups is sensitive to which of BA.1, BA.2, BA.5, or XBB.1.5 the serum group has been exposed (Figure 5B). In the serum groups from China, titers are higher for variants nearer to BA.2 in the antigenic map (BA.2, BA.5, and all XBB.x variants) in the BA.2 + BA.5/BF.7 serum group than in the BA.1 + BA.5/BF.7 serum group - and vice versa, with higher titers in the BA.1 infected serum group for variants nearer to BA.1 (Figure 5C). This effect is also reflected in the position of the peak of the antibody landscapes for these serum groups (Figure 5B). For the XBB.1.5 infection serum group from Denmark, titers to XBB.x variants are higher than seen in the BA.5 infection cohort, and the peak of the landscape is shifted further in the XBB.x direction.

This shift can explain why titers to XBB.1.5 and XBB.1.16 are higher than titers to BA.2.86 in serum groups from China with two BA.x infections, despite not having had an exposure to XBB.x - because repeated exposure to BA.x in these groups has moved the peak of the landscape to a region of antigenic space that is nearer to the XBB.x variants than to BA.2.86.

### Vaccination with XBB.1.5 raises and broadens the antibody landscape but does not shift the peak

Vaccination with XBB.1.5 in both the ADARC (USA) and Moderna/Duke (USA) datasets increases titers to all variants relative to pre-XBB.1.5-vaccination serum groups (Figure 6). In all cohorts, fold increase in titer after XBB.1.5 vaccination is greater for XBB.x variants and BA.2.86/JN.1 than for less antigenically advanced variants (i.e. variants which are less antigenically distant from the Anc./D614G variant). In cohorts with no confirmed prior infection, XBB.1.5 vaccination results in a 12.1-13.3x fold increase in BA.2.86/JN.1 titer, compared to a fold increase of 6.8-11.2x to BA.5 (Figure 6). The larger fold increase in titer for more antigenically advanced variants is reflected in a flattening of the antibody landscape in the post-vaccination serum groups relative to the corresponding pre-XBB.1.5-vaccination serum groups (Figure 6), with the peak of the landscape remaining near to D614G.

**Figure 6:**
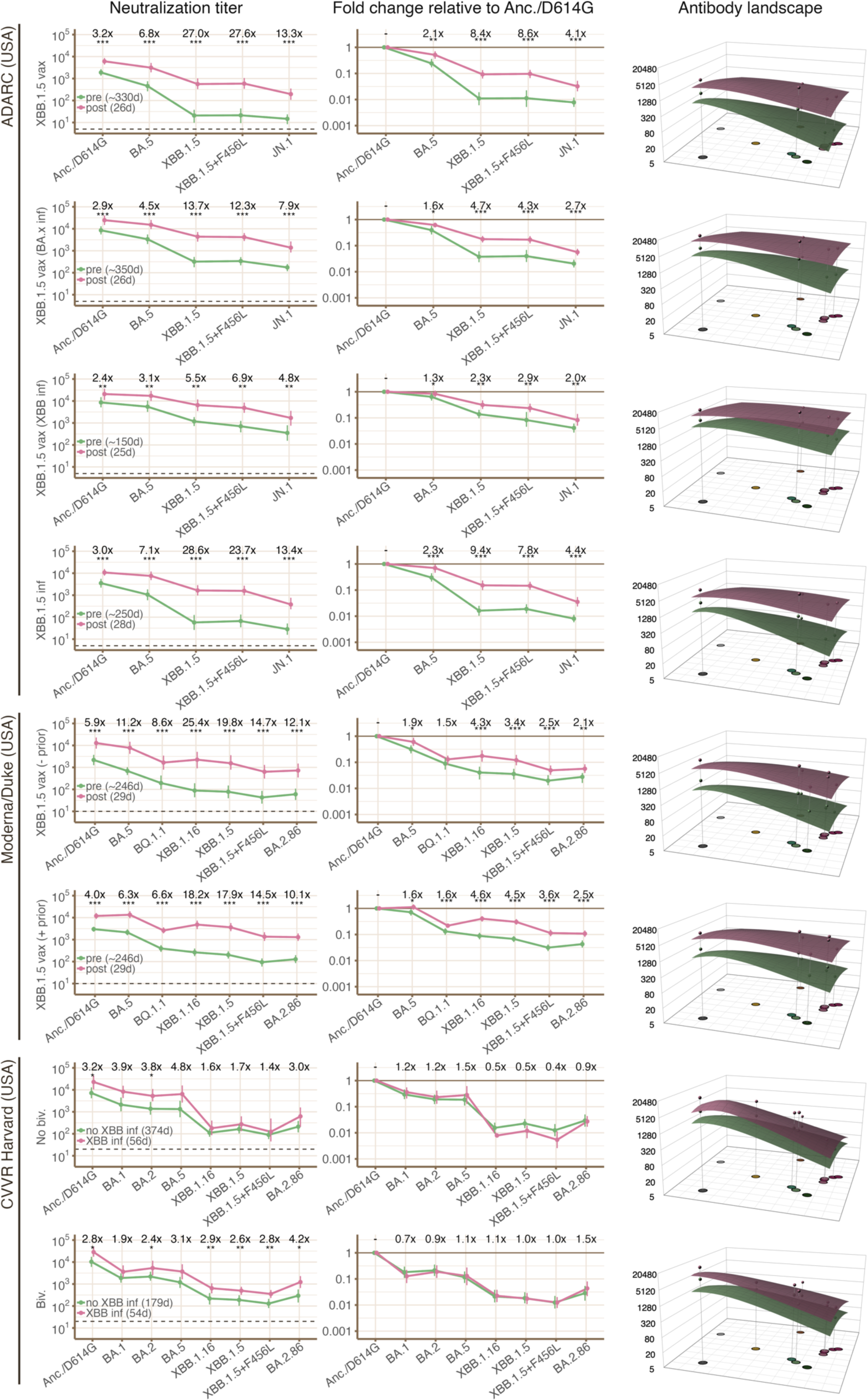
Geometric mean neutralization titers (left column), geometric mean fold change relative to the Anc./D614G variant (center column), and antibody landscapes (right column) are shown for selected pairs of serum groups from individual laboratories which allow direct comparison of the effect of XBB vaccination status or infection status. For each pair, fold changes from unexposed to exposed serum groups (either by infection or vaccination) are written in the “neutralization titer” column. In the “Fold change relative to Anc./D614G” column, differences in fold change relative to Anc./D614G between the unexposed and exposed serum group in each pair are written. Asterisks indicate level of statistical significance (* = p<0.05; ** = p<0.01; *** = p<0.001) as determined by the Wilcoxon paired difference test for paired data (from ADARC (USA) and Moderna/Duke (USA)), or by the Wilcoxon signed rank test for unpaired data (from CVVR Harvard (USA)). 95% confidence intervals are shown, calculated by the non-parametric bootstrap using 1000 replicates. The number of days between the previous exposure (vaccination or infection) and the sample is given in parentheses in the legend for each serum group when known. Abbreviations are used in serum group names and exposure history: “vax” = vaccinated; “inf” = infected; “Anc.” = ancestral/Wuhan-1 SARS-CoV-2 virus; “biv.” = bivalent.

While fold increase is larger for BA.2.86/JN.1 than for BA.x and Anc./D614G variants, fold increase is largest for XBB.x variants, and is significantly larger for XBB.1.5 than for BA.2.86/JN.1 in all four XBB.1.5 vaccination datasets which have not had confirmed prior XBB infection (Figure S1). As XBB.1.5 exposure causes a larger increase in titer to XBB.1.5 than to BA.2.86, all serum groups with previous XBB.1.5 exposure from ADARC (USA) have lower estimated BA.2.86 titers than XBB.1.15 titers (Figure 3C, S9). In the datasets from Moderna/Duke (USA), XBB.1.5 titers are higher than BA.2.86 titers both pre and post XBB.1.5 vaccination, with the XBB.1.5/BA.2.86 titer ratio increasing after vaccination due to the greater boosting of XBB.1.5 titers (Figure 3C, S10).

### XBB infection data suggests that flattening of the antibody landscape may be transient

The XBB.1.5 infection cohort from ADARC (USA) shows a pattern of raising and flattening the antibody landscape, similar to that seen in XBB.1.5 vaccination cohorts from ADARC (USA) and Moderna/Duke (USA) (Figure 6). However, in the XBB.x infection cohorts from CVVR Harvard, the increase in neutralization titer is similar against all variants compared to the non-infected cohort (Figure 6). As such, the antibody landscape is raised but not flattened after XBB.x infection. In addition to confounds such as different assays and prior history of vaccination and infections, this difference may be caused by the shorter time since exposure for the XBB.1.5 infection and post-vaccination serum groups from ADARC (USA) and Moderna (USA) (<30 days) than post infection serum groups from CVVR Harvard (USA) (54 and 56 days). This suggests that the flattening of the antibody landscape after XBB.x exposure may be transient, and that titers to more antigenically advanced variants may wane faster than those to earlier variants, eventually resulting in similar fold increase between pre and post vaccination titers for all variants, as seen in data from CVVR Harvard. A longitudinal sampling from the same cohort/trial will be needed for a definitive conclusion. Previous studies have observed faster waning of titers against Omicron variants than against Anc./D614G after bivalent vaccination (Lasrado, Collier, Miller, et al. 2023; Jacobsen et al. 2023; Chalkias et al. 2022),.

### BA.x infection and vaccination are less effective at raising the antibody landscape in the UK and USA

Infection or bivalent vaccination with a BA.x variant in cohorts from the UK or USA had a smaller effect on neutralization titers to XBB.x and BA.2.86 variants than XBB.1.5 infection or vaccination (Figures 2, S2). In serum groups which have received a bivalent vaccine <30 days ago, fold increase to XBB.1.5 is 3.1-5.4x, and to BA.2.86 is 1.6-4.0x. These fold increases decline in >150 day post-vaccination serum groups, to 1.5-2.9x against XBB.1.5, and no significant fold increase against BA.2.86 (1.0-1.2x). Two BA.x infections in the PKU (China) serum groups cause a larger increase in XBB.x and BA.2.86 titers compared to single BA.x infection cohorts, of 8.9-11.0x against XBB.1.5, and 3.3-5.4x against BA.2.86.

In data from China, titers to BA.2.86 are similar between double infection serum groups with BA.x + XBB.x infection and those with two BA.x infections, and titers to BA.2.86 are in fact slightly lower in the single infection XBB.x serum group than the two BA.x infection serum groups (Figure 2). Titers to BA.2.86 are also similar between the XBB.1.5 and the BA.5 infection serum groups from SSI (Denmark) (Figure 2).

### Do any serum groups see BA.2.86 as particularly antigenically distant relative to pre-BA.2.86 variants?

In addition to identifying the serum groups which produce lower titers to BA.2.86 than to XBB.1.5/16, we identified serum groups with low BA.2.86 titers relative to an average across all pre-BA.2.86 variants (see Methods for details) (Figure 4B). There is generally good agreement in BA.2.86’s antibody escape for most serum groups - fold reduction relative to the average across earlier variants is 3.2-7.9x in 24 of 31 serum groups (with titrations against a sufficient number of variants).

Four serum groups have a fold reduction in titer from pre-BA.2.86 variants to BA.2.86 of greater than 7.9x: the two pre-XBB.1.5-vaccination serum groups from Moderna/Duke (USA) with fold reductions of 8.8x and 9.2x; the Elderly post-bivalent vaccination cohort from MRC-CVR (UK) (13.2x fold reduction); and the post-bivalent cohort from VDEC (UK) (9.0x fold reduction) (Figure 4B). The two post-bivalent serum groups from MRC-CVR (UK) and VDEC (UK) are the only two serum groups with a >64x fold reduction from Anc./D614G to BA.2.86, and the only serum groups with lower BA.2.86 titers than XBB.1.5 titers outside of the four main groups identified earlier (Figures 3C, S3). This result contrasts with post-bivalent vaccination serum groups from the USA - a potential explanation is that bivalent booster doses in the UK were offered only to individuals with risk factors, including age, which may bias the cohort to an older age group.

Of the three serum groups with a fold reduction to BA.2.86 of less than 3.2x, two are post-XBB.1.5 vaccination serum groups with confirmed prior BA.x or XBB.1.5 infections. The remaining serum group is from MRC-CVR (UK), where the lower limit of detection is higher than in data from other laboratories - the full extent of escape for more recent variants relative to Anc./D614G and BA.x is therefore not fully captured, with most titers to XBB and BA.2.86 being below the limit of detection (Figure S11).

## Discussion

In summary, we find that there are differences in the neutralization profiles of cohorts from countries with different exposure histories, which affect the relative extent of antibody escape for BA.2.86 compared to other cocirculating variants.

In the UK and USA, antibody landscapes remain centered on the Anc./D614G variant regardless of infection or vaccination history. This suggests that the antibody response is heavily imprinted on Anc./D614G-like variants, with subsequent exposures boosting the cross-reactive component of the response without generating a novel component of the response. Despite this, vaccination with XBB.1.5 is effective at raising and broadening the antibody landscape, with titers increasing most substantially to antigenically advanced variants. This includes a large increase in titers against BA.2.86, which is much larger than the increase in BA.2.86 titers after bivalent BA.x vaccination, despite BA.2.86 being a similar antigenic distance from XBB.x as from BA.x. This suggests that vaccination with an antigenically advanced variant may be more broadly immunogenic than vaccination with an older variant when there is strong imprinting on earlier variants, therefore providing greater antibody-mediated protection even against variants which are substantially antigenically diverged from the variant included in the vaccine.

In serum groups from China, two BA.x infections result in a shift of the peak of the antibody landscape towards the BA.x region of antigenic space, resulting in higher titers to XBB than to BA.2.86 due to the greater proximity of BA.x to XBB than to BA.2.86 in the antigenic map. Further, serum groups with infections by BA.x have similar (for double infection serum groups) or slightly higher (for single infection serum groups) titers to BA.2.86 than those with XBB.x infections. Together, these suggest weaker imprinting on Anc./D614G-like variants than in the UK and USA, allowing a novel component of the antibody response to be generated that is focused on more recently encountered variants, even during exposure to less antigenically divergent variants such as BA.x. A number of factors may cause this weaker imprinting on Anc./D614G in China, including (1) lower immunogenicity of the inactivated virus vaccines compared to mRNA vaccines used in the UK, USA, and Denmark; (2) a smaller number of vaccinations in China (three in China, compared to up to six in the UK/USA when including bivalent boosters); and (3) a smaller number of infections with pre-Omicron variants, due to particularly strong restrictions on the spread of the virus early in the pandemic.

Similarly, serum groups from Denmark with BA.x or XBB.1.5 infections also showed a shifted peak of the antibody landscape, and similar titers to BA.2.86. In this case, the differing epidemiological history of Denmark may explain the ability to overcome imprinting on the Anc./D614G variant. Denmark experienced a smaller number of pre-Omicron infections per individual than the UK or USA, with a substantially lower rate of test positivity in Denmark than the UK or USA during the period of circulation of pre-Omicron variants, and a slightly higher rate during the period of early Omicron variant circulation (Figure 1C, Figure S4). The lower number of pre-Omicron variant infections in Denmark may have resulted in weaker imprinting on pre-Omicron variants compared to the UK and USA, with the large number of subsequent Omicron infections able to overcome the imprinting, analogous to the effect of two BA.x infections in serum groups from China.

The primary limitation of this study is that a number of comparisons are made across different cohorts, which additionally are characterized using different neutralization assay protocols across laboratories (Table S1). It is possible that differences in assay protocol result in systematic differences in measured neutralization titers, limiting comparability across the laboratories. Side-by-side characterization of serum samples collected from a range of countries would help to confirm that the measured differences are caused by differences in population immunity rather than by differences in assay protocol. Further, conclusions about the effect of exposure to variants on the immune profile are often made by comparing neutralization profiles across different cohorts. This is confounded by the possibility that pre-exposure immune profiles differ between the cohorts, even if they are measured in the same laboratory. Conclusions about the differences in exposure history which may have caused the observed differences between cohorts in this study could be strengthened by comparing appropriate samples from longitudinal cohorts.

In our cohorts, the relative escape of XBB.1.5 and BA.2.86 from antibody-mediated immunity varied between countries and between exposure histories, demonstrating how the fitness of a variant and therefore the threat it poses may vary geographically. As such, it is especially important to consider antigenicity with respect to sera from a wide variety of locations and exposure histories when trying to judge the likely future epidemiological trajectory of a variant. With the subsequent substantial circulation of the JN.1 subvariant, it would be interesting to determine whether there is variation in the degree of antibody escape between different exposure histories for this variant. JN.1 possesses only a single additional spike substitution relative to BA.2.86, compared with the large number of substitutions that define BA.2.86.

Differences in the response to exposure between countries may be also relevant for decisions about vaccine antigen composition. In sera from the UK and USA, infection or vaccination with XBB.x is consistently effective at increasing titers to BA.2.86, while exposure to BA.x has a smaller and more variable effect, which may decline further as the time since exposure increases. This suggests that the policy of using XBB.1.5-based vaccine boosts in these regions is sound not only for protection against XBB.1.5, but also BA.2.86 and other antigenically advanced variants that may arise in the future, as the more recent XBB.1.5 variants evade the substantial imprinting on earlier variants to allow greater boosting of the antibody response. Contrastingly, BA.2.86 titers are not boosted more effectively by XBB.x infection relative to BA.x infection in sera from China or Denmark, although direct comparison of vaccination cohorts is not possible for these cohorts. Though there is not sufficient data to make definitive statements about optimal vaccine composition, the early infection and vaccination history and therefore the focus and strength of imprinting may be important to consider when making this decision in different countries.

## Methods

### Individual group methodologies

#### PKU (China) - Peking University

Whole blood samples were diluted 1:1 using phosphate buffered saline (PBS) (Invitrogen, C10010500BT) with 2% fetal bovine serum (FBS) (Hyclone, SH30406.05) and then underwent Ficoll (Cytiva, 17-1440-03) density gradient centrifugation for separation of plasma and peripheral blood mononuclear cells (PBMCs). The plasma was retrieved from the top layer post-centrifugation. Plasma samples were aliquoted and stored at −80°C.

Vesicular stomatitis virus (VSV) pseudotyped virus was constructed as described before (Yisimayi et al. 2024). Plasma samples were diluted in DMEM (Hyclone, SH30243.01) and mixed with the pseudovirus. This mixture was incubated in a 96-well plate at 37°C and 5% CO2 for an hour. Huh-7 cells (JCRB, 0403) were then added to the mixture and incubated for 24 hours. Subsequently, half of the supernatant was removed, and a luciferase assay was performed using Bright-Lite Luciferase Assay Substrate and Buffer (Vazyme, DD1209-03-AB). Luminescence was measured using a microplate spectrophotometer (PerkinElmer, HH3400). The IC50 value was calculated using a four-parameter logistic regression model in PRISM (version 9.0.1).

#### SSI (Denmark) - Statens Serum Institut

Virus neutralization titers were determined using a live virus microneutralization test with SARS-CoV-2 clinical isolates (Frische et al. 2022). In brief, a 2-fold serial dilution of heat-inactivated serum/plasma samples was pre-incubated with 300 × 50% tissue culture infectious dose SARS-CoV-2 virus and subsequently transferred to a Vero E6 cell monolayer in a 96-well tissue culture plate. The following day, the cells were fixed with acetone and the level of virus infection was measured in a standard ELISA targeting the SARS-CoV-2 nucleocapsid protein. The 50% virus neutralization titer was determined for each sample using a four-parameter logistic regression curve fit with the cut-off calculated from virus- and mock-infected control wells on the corresponding assay plate. The SARS-CoV-2 isolate sequences are available in the European Nucleotide Archive (project number PRJEB67449).

#### MRC-CVR (UK) - MRC University of Glasgow Centre for Virus Research

HEK293, HEK293T and 293-ACE2 cells were maintained in Dulbecco’s modified Eagle’s medium (DMEM) supplemented with 10% fetal bovine serum, 200 mM L-glutamine, 100 μg/ml streptomycin and 100 IU/ml penicillin. HEK293T cells were transfected with the appropriate SARS-CoV-2 spike gene expression vector in conjunction with lentiviral vectors p8.91 and pCSFLW using polyethylenimine (PEI, Polysciences, Warrington, USA). Pseudotype-containing supernatants were harvested 48 hours post-transfection, aliquoted and frozen at −80C prior to use. The SARS-CoV-2 spike glycoprotein expression constructs for ancestral B.1 and Omicron have been described previously (Willett et al. 2022). 293-ACE2 target cells were maintained in complete DMEM supplemented with 2 μg/ml puromycin.

Neutralizing activity in each sample was measured by a serial dilution approach. Each sample was serially diluted in triplicate from 1:50 to 1:36,450 in complete DMEM prior to incubation with approximately 1 x 10^6^ CPS per well of HIV (SARS-CoV-2) pseudotypes, incubated for 1 h, and plated onto 293-ACE2 target cells. Luciferase activity was quantified after 48–72 h by the addition of Steadylite Plus chemiluminescence substrate and analysis on a Perkin Elmer EnSight multimode plate reader (Perkin Elmer, Beaconsfield, UK). Antibody titre was then estimated by interpolating the point at which infectivity had been reduced to 50% of the value for the ‘no serum’ control samples.

#### VDEC (UK) - Vaccine Development and Evaluation Centre

As described in (Coombes, Bewley, Le Duff, Alami-Rahmouni, et al. 2023), viruses were isolated from sequence-confirmed clinical swabs at UKHSA. SARS-CoV-2 variants were isolated and propagated on Vero/hSLAM cells (ECACC 04091501) before being subjected to quality control checks as previously described ((Coombes, Bewley, Le Duff, Hurley, et al. 2023). Briefly, titrations were performed using a focus-forming assay with immunostaining for nucleocapsid to visualize foci (Ryan et al. 2023). Sighting for each variant into the Focus Reduction neutralization Test (FRNT) to calculate dilution at which a median of 130 foci could be counted in non-neutralized control wells was performed. Median neutralizing antibody titers (ND50) were measured with FRNT and calculated with a Probit regression analysis using R v4.1.2 (R Core Team, Vienna, Austria) as described previously (Bewley et al. 2021; Coombes, Bewley, Le Duff, Hurley, et al. 2023).

#### Pirbright/UKHSA (UK) - The Pirbright Institute / UK Health Security Agency

Testing was performed by Pirbright-UK with samples from the UKHSA-CONSENSUS study, which is a prospective longitudinal audit of antibody levels and T cell responses in vaccinated adults.

The method used to determine neutralization titers from CONSENSUS study sera is described in detail in (Newman et al. 2022). Briefly, SARS-CoV-2 lentiviral pseudotypes were generated by transfection of HEK293T cells with plasmids expressing SARS-CoV-2 spike variants, p8.91 (encoding HIV-1 *gag-pol*) and CSFLW (encoding a firefly luciferase reporter gene). After harvest and titration of pseudovirus, a dilution equivalent to 10^5^-10^6^ signal luciferase units was mixed with three-fold serial dilutions (1/30-1/21870) of heat-inactivated sera in 96-well white tissue-culture plates in a final volume of 100µl and incubated at 37°C for 1 hour. Target cells (HEK293T-hACE2) were added at a density of 2×10^4^ cells in 100µl to each well and then incubated at 37°C for 48 h. Firefly luciferase activity was measured after replacing the media with 50µl Bright-Glo luciferase reagent and measuring luminescence on a GloMax-Multi^Tm^ Detection system (Promega). Neutralization titers were calculated by interpolating the reciprocal serum dilution at which there was a 50% inhibition of luciferase activity relative to untreated controls (ND_50_).

#### CVVR Harvard (USA) - Center for Virology and Vaccine Research

Neutralizing antibody (NAb) titers against SARS-CoV-2 variants were determined using pseudotyped viruses expressing a luciferase reporter gene, as previously described (Lasrado, Collier, Hachmann, et al. 2023). In brief, HEK293T-hACE2 cells were seeded in 96-well tissue culture plates at a density of 2 × 10^4^ cells per well overnight. Three-fold serial dilutions of heat-inactivated serum samples were prepared and mixed with 60 μl of pseudotyped SARS-CoV-2 variant viruses and incubated at 37 °C for 1 h before adding to HEK293T-hACE2 cells. 48 h later, cells were lysed in Steady-Glo Luciferase Assay (Promega) according to the manufacturer’s instructions. SARS-CoV-2 neutralization titers were defined as the sample dilution at which a 50% reduction (NT50) in relative light units was observed relative to the average of the virus control wells.

#### ADARC (USA) - Aaron Diamond AIDS Research Center

Serum neutralizing antibody titers against SARS-CoV-2 variants were determined on VSVΔG pseudotyped viruses expressing a luciferase reporter gene, as previously described (DOI: 10.1038/s41586-020-2571-7). The viral titer of each variant was titrated and normalized before neutralization assays. Serum samples were diluted in triplicate in 96-well plates, starting from a 12.5-fold dilution, and then incubated with an equal volume of virus for 1 h at 37°C before adding 4°10^4^ cells/well of Vero-E6 cells. The cells were then cultured overnight, harvested, and lysed for luciferase activity measurement using SoftMax Pro v.7.0.2 (Molecular Devices). Reductions in luciferase activity corresponding to various serum dilutions were determined. ID_50_ titers, reflecting a 50% reduction in luciferase activity, were calculated by fitting the data to a non-linear five-parameter dose-response curve using GraphPad Prism V.10.

#### Moderna/Duke (USA) – Moderna & Duke University Medical Center

The lentivirus-based SARS-CoV-2 spike-pseudotyped virus neutralization assay performed at Duke quantifies nAbs as a reduction in firefly luciferase expression after a single round of infection in 293T-ACE2 cells (Gilbert et al. 2022). Serial dilution of serum samples was used to produce a dose-response curve. Neutralization was measured as the serum dilution at which relative luminescence units (RLU) was reduced by 50% (ID50) relative to mean RLU in virus control wells (cells + virus but no sample) after subtraction of mean RLU in cell control wells (cells only).

### Epidemiological data

Epidemiological data about the number of BA.2.86 cases and BA.1/2/4/5 cases globally and per-country for Figures 1A and 1B was obtained from covSPECTRUM (Chen et al. 2022) using the query “BA.2.86*”. Data for the number of confirmed infections for Figure 1C and about test positivity rates were obtained from Our World in Data (Edouard Mathieu, Hannah Ritchie, Lucas Rodés-Guirao, Cameron Appel, Charlie Giattino, Joe Hasell, Bobbie Macdonald, Saloni Dattani, Diana Beltekian, Esteban Ortiz-Ospina and Max Roser n.d.) which uses data from the WHO (World Health Organization n.d.). To estimate the number of cases of Omicron or pre-Omicron infection in Figures 1C and S4, the number of confirmed infections or test positivity rate on a particular date was multiplied by the proportion of sequences which were descendants of B.1.1.529 (Omicron) which were sampled within 1 week of that date.

### Antigenic map

The antigenic map was constructed using antigenic cartography (Smith et al. 2004) with the “Racmacs” package (Sam Wilks 2023) in R (v4.2.1 (R Core Team 2022)) using neutralization data from PKU (China) using 1 week post immunization mouse serum. As described in (Rössler et al. 2023), the table of neutralization titers is converted into a distance table by calculating the log2 fold change from the maximum titer for each serum to all other titers per serum. Coordinates for each serum and variant pair are then optimized such that their Euclidean distance in the map matches their table distance. A detailed description of the algorithm is given by Smith et al. and the reference page of the Racmacs package (Smith et al. 2004; Sam Wilks 2023).

### Antibody landscapes

Antibody landscapes were constructed using the “ablandscapes” package (Samuel Wilks 2021) in R for all serum groups where there were titrations against at least the D614G virus, one of BA.1/2/4/5, one XBB.x variant, and BA.2.86. For each serum group, single-cone landscapes were fit to each individual serum, and the GMT landscape calculated by taking the average of the landscape for the individual sera in that group.

### Calculating Geometric Mean Titers (GMTs)

When calculating GMTs for a serum group against a variant, titers below the lower limit of detection are treated as 0.5x the lower limit of detection.

### Estimating BA.2.86 titers from JN.1 titers

In the eight pre- and post-XBB.1.5 vaccination serum groups from ADARC, titrations were not performed against BA.2.86, but instead against JN.1. As JN.1 is not included in the antigenic map used to fit the antibody landscapes and is not included in titration sets from most other laboratories, we estimate BA.2.86 titer values for each serum in these serum groups by multiplying the titer against JN.1 by 3x. The value of 3x was chosen by considering the fold reduction from BA.2.86 to JN.1 from three studies: 2.1x, found in the BA.5/BF.7 + XBB infection cohort from PKU (China) (Yang et al. 2024); 3.8-4.5x, found in a study of sera collected in Japan (Kaku et al. 2024); and 3.4x, found in VDEC’s study of sera from individuals vaccinated with 3x ancestral vaccines and a BA.1 bivalent booster. The estimated titers are used to identify serum groups with low BA.2.86 titers relative to other serum groups for discussion, and to fit antibody landscapes for visualization. The estimated titers are not included in any reported statistics.

### Mean across pre-BA.2.86 variants

The GMT across pre-BA.2.86 variants for Figure 4B was calculated for each serum group by first taking the geometric mean of the GMTs against BA.1, BA.2, and BA.5 to produce a “BA.x GMT”, and the geometric mean of the GMTs against XBB.1.5 and XBB.1.16 to produce an “XBB.1.5/16 GMT“. If a subset of these variants was not present, a mean over those which were was taken to produce the BA.x mean and the XBB.1.5/16 mean. The mean across pre-BA.2.86 variants for a serum group is then the geometric mean of the Anc./D614G GMT, the BA.x GMT, and the XBB.1.5/16 GMT.

### Grouping of variants

Different laboratories sometimes performed titrations against different subtypes of variants, with slight differences in spike sequence, as shown in the table below. Figure S5 compares titers for each pair of variants in the table below in cases where a single serum is titrated against more than one variant, with titers typically varying by less than a 2-fold dilution in each case. As such, in cases where additional substitutions were not contained in the RBD, additional substitutions were ignored and matched to another variant as in the table below.

| Variant name in this paper | Variants included |
| --- | --- |
| Anc./D614G | Wuhan-1/Ancestral, D614G |
| XBB(.1) | XBB, XBB.1 (= XBB+G252V) |
| XBB.1.5+F456L | EG.5 (XBB.1.5+F456L), EG.5.1 (XBB.1.5+Q52H+F456L) |
| BA.2.86 | BA.2.86, BA.2.86+I670V |

### Defining serum groups with high BA.2.86 escape from neutralization relative to historic variants

We identified serum groups with low BA.2.86 titers relative to each of Anc./D614G, BA.x, and XBB.1.5/16. To do this, we calculated the fold change in titer from the Anc./D614G GMT, the BA.x GMT, and the XBB.1.5/16 GMT (the latter two means are defined in the “Mean across pre-BA.2.86 variants’’ section of Methods). High fold change relative to XBB.1.5/16 is defined as any fold reduction greater than 1x, relative to BA.x as a fold reduction greater than 16x, and relative to Anc./D614G as a fold reduction greater than 64.

### Statistical analyses

Null Hypothesis Significance Testing on paired data was performed using the Wilcoxon matched pairs test, and on unpaired data using the Wilcoxon rank sum test. Confidence intervals were calculated using the non-parametric bootstrap with 1000 replicates.

## Acknowledgements

The authors thank Yann Le Duff, Nassim Alami-Rahmouni and Sarah Kempster from MHRA for collecting and sharing samples included in the serum panels analyzed at UKHSA.

The authors thank the DOVE Consortium for sera analyzed by MRC-CVR. The DOVE Consortium includes Brian J Willett, Nicola Logan, Sam Scott, Chris Davis, Therese McSorley, Patawee Asamaphan, Margaret J Hosi, Paula Olmo, Joe Grove, Richard Orton, Antonia Ho, John Haughney, David L Robertson, and Emma C Thompson.

---

**Figure S1.**
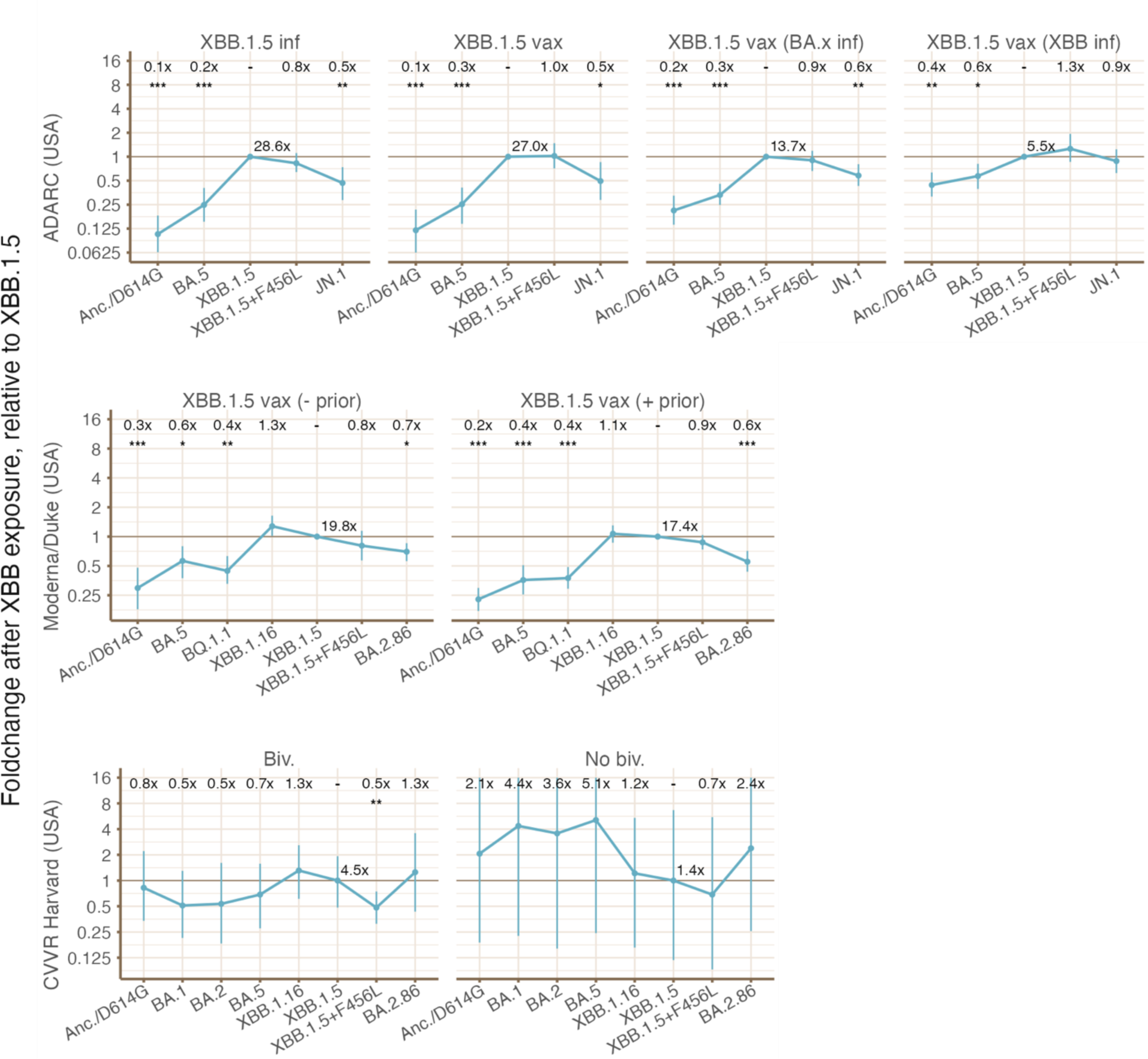
Fold change between the unexposed to the exposed serum group is compared for each variant against XBB.1.5, for each pair of serum groups shown in Figure 6 (such that a variant which shows a larger fold increase in titer than XBB.1.5 would have a value of greater than one in this figure). The fold increase in XBB.1.5 titer between pre- and post-vaccination serum groups is written next to the XBB.1.5 data point. For data from ADARC (USA) and Moderna/Duke (USA), fold increase is 0.5-0.9x smaller for BA.2.86 than XBB.1.5. Asterisks indicate level of statistical significance (* = p<0.05; ** = p<0.01; *** = p<0.001) as determined by the Wilcoxon paired difference test for paired data (from ADARC (USA) and Moderna/Duke (USA)), or by the Wilcoxon signed rank test for unpaired data (from CVVR Harvard (USA)). 95% confidence intervals are shown, calculated by the non-parametric bootstrap using 1000 replicates. Abbreviations are used in serum group names and exposure history: “vax” = vaccinated; “inf” = infected; “biv.” = bivalent.

**Figure S2.**
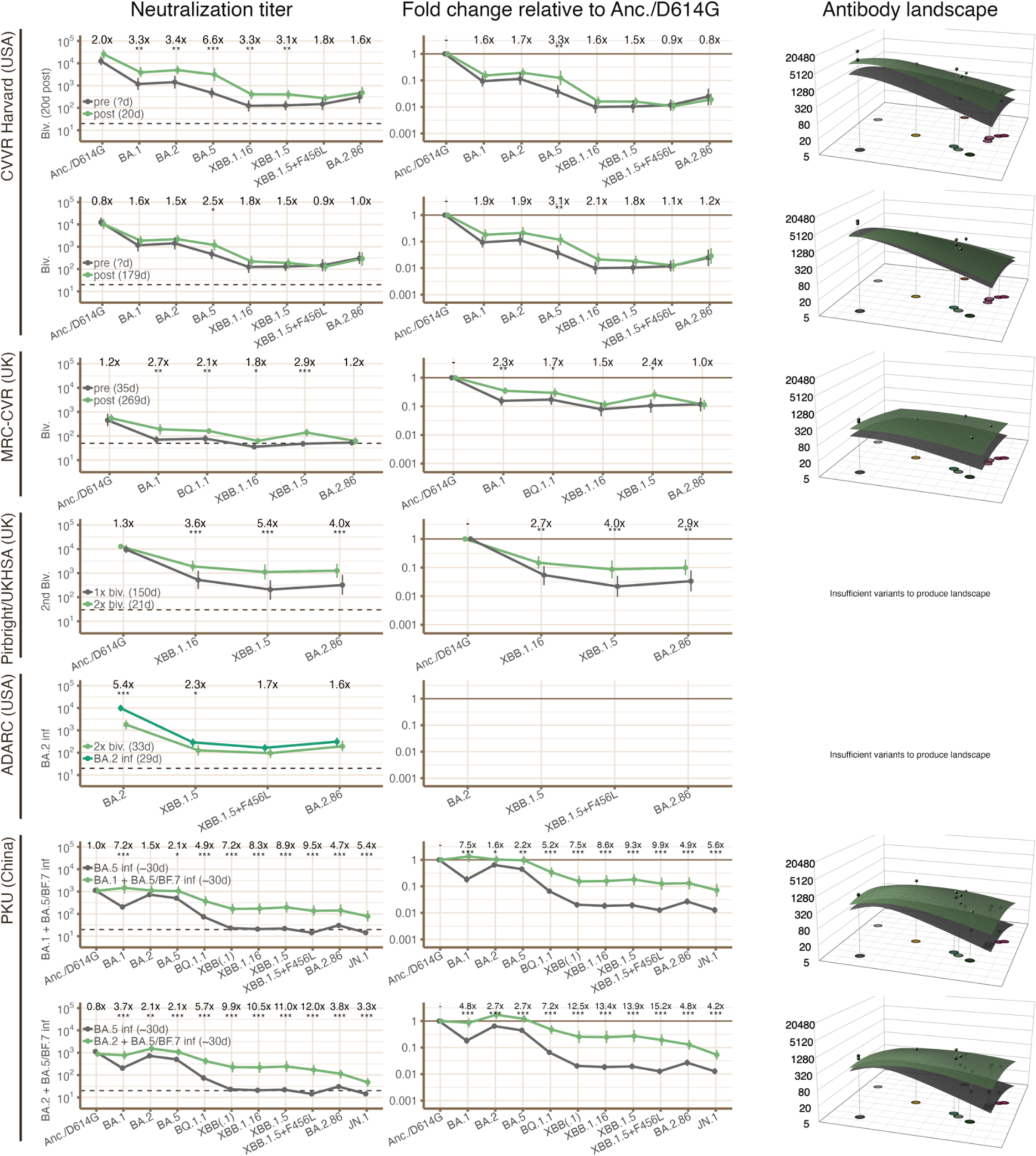
Geometric mean neutralization titers (left column), geometric mean fold change relative to the Anc./D614G variant (center column), and antibody landscapes (right column) are shown for selected pairs of serum groups from individual laboratories which allow direct comparison of the effect of BA.x vaccination status or infection status. For each pair, fold changes from unexposed to exposed serum groups (either by infection or vaccination) are written in the “neutralization titer” column. In the “Fold change relative to Anc./D614G” column, differences in fold change relative to Anc./D614G between the unexposed and exposed serum group in each pair are written. Asterisks indicate level of statistical significance (* = p<0.05; ** = p<0.01; *** = p<0.001) as determined by the Wilcoxon paired difference test for paired data (from Pirbright/UKHSA (UK)), or by the Wilcoxon signed rank test for unpaired data (from all other labs in this figure). 95% confidence intervals are shown, calculated by the non-parametric bootstrap using 1000 replicates. The number of days between the previous exposure (vaccination or infection) and the sample is given in parentheses in the legend for each serum group when known. Abbreviations are used in serum group names and exposure history: “vax” = vaccinated; “inf” = infected; “Anc.” = ancestral/Wuhan-1 SARS-CoV-2 virus; “biv.” = bivalent.

**Figure S3.**
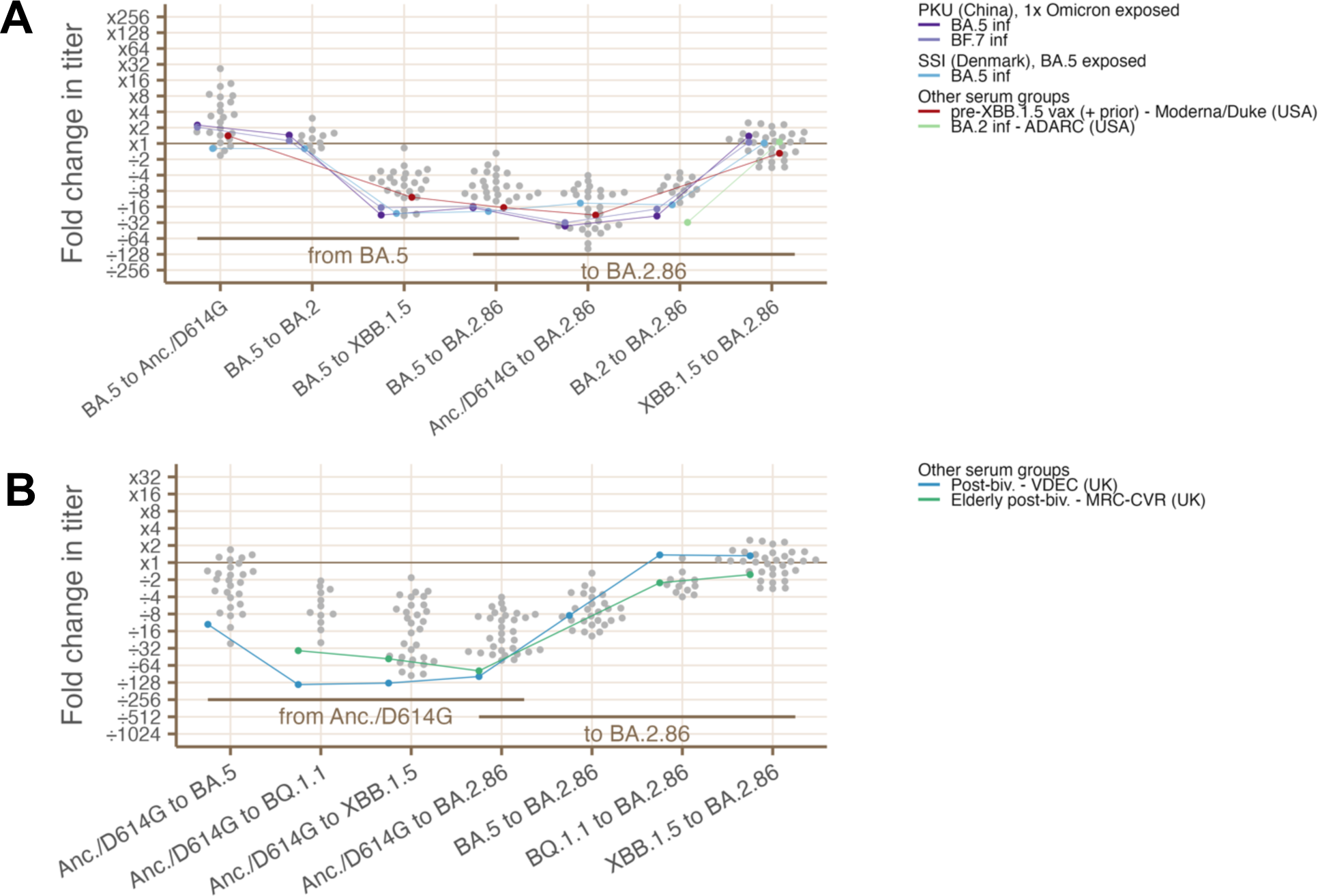
**(A)** Comparisons of fold change in neutralization titer for serum groups which show reductions in neutralization titer of >16x against BA.2.86 relative to BA.x (see Methods for details of how these serum groups are identified). Fold change values are shown from BA.5 to each other variant, and then from each other variant to BA.2.86. Serum groups showing a >16x reduction titer from BA.2.86 to BA.x are plotted in color, compared to all other serum groups which are plotted in gray. (**B**) As in **A**, highlighting serum groups showing a >64x reduction in neutralization titer from Anc./D614G to BA.2.86. Abbreviations are used in serum group names and exposure history: “vax” = vaccinated; “inf” = infected; “biv.” = bivalent.

**Figure S4.**
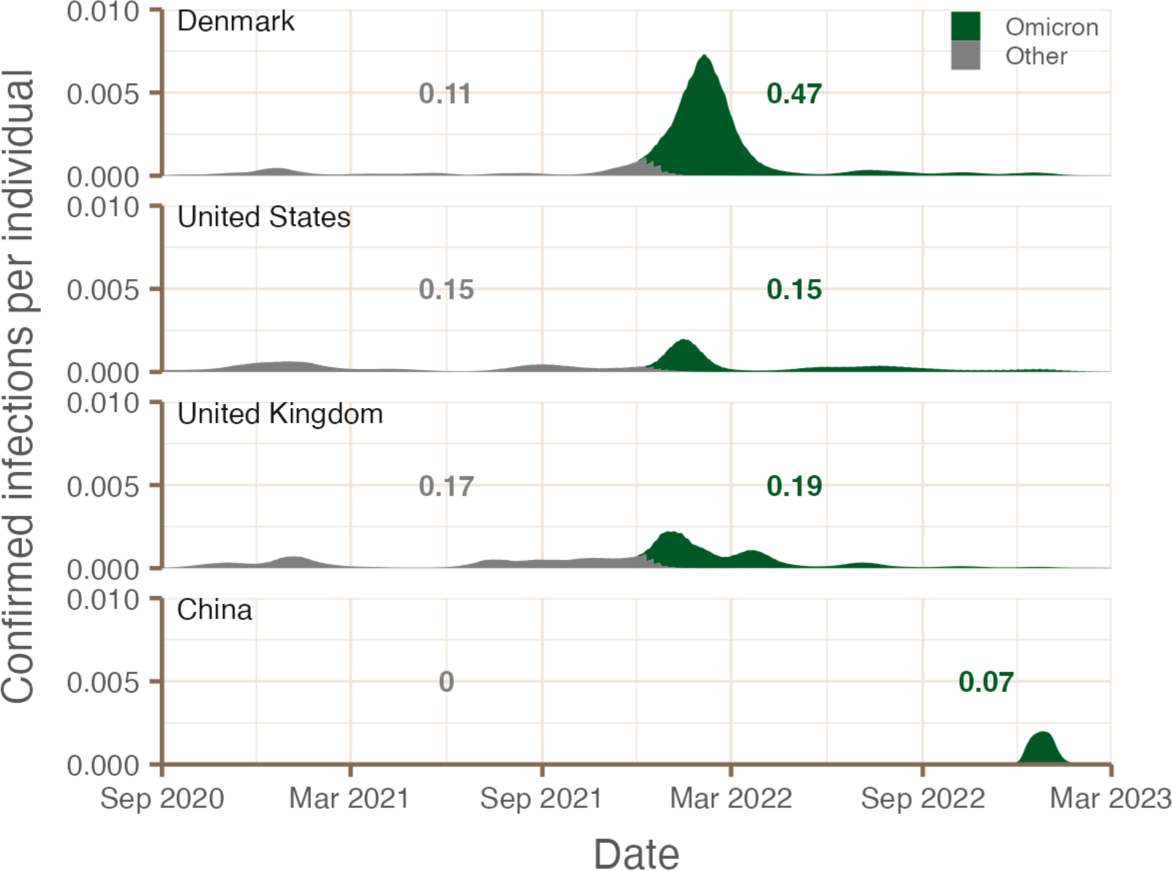
Equivalent to Figure 1C, calculated using number of confirmed infections instead of test positivity. The test positivity rates suggest a substantially smaller wave of pre-Omicron infections followed by a slightly larger BA.x wave in Denmark than in the USA and UK, while counts of confirmed infections suggest a slightly smaller wave of pre-Omicron infections followed by a substantially larger wave of Omicron infections. This difference is due to the higher rate of testing in Denmark compared to the UK and USA early in the pandemic.

**Figure S5.**
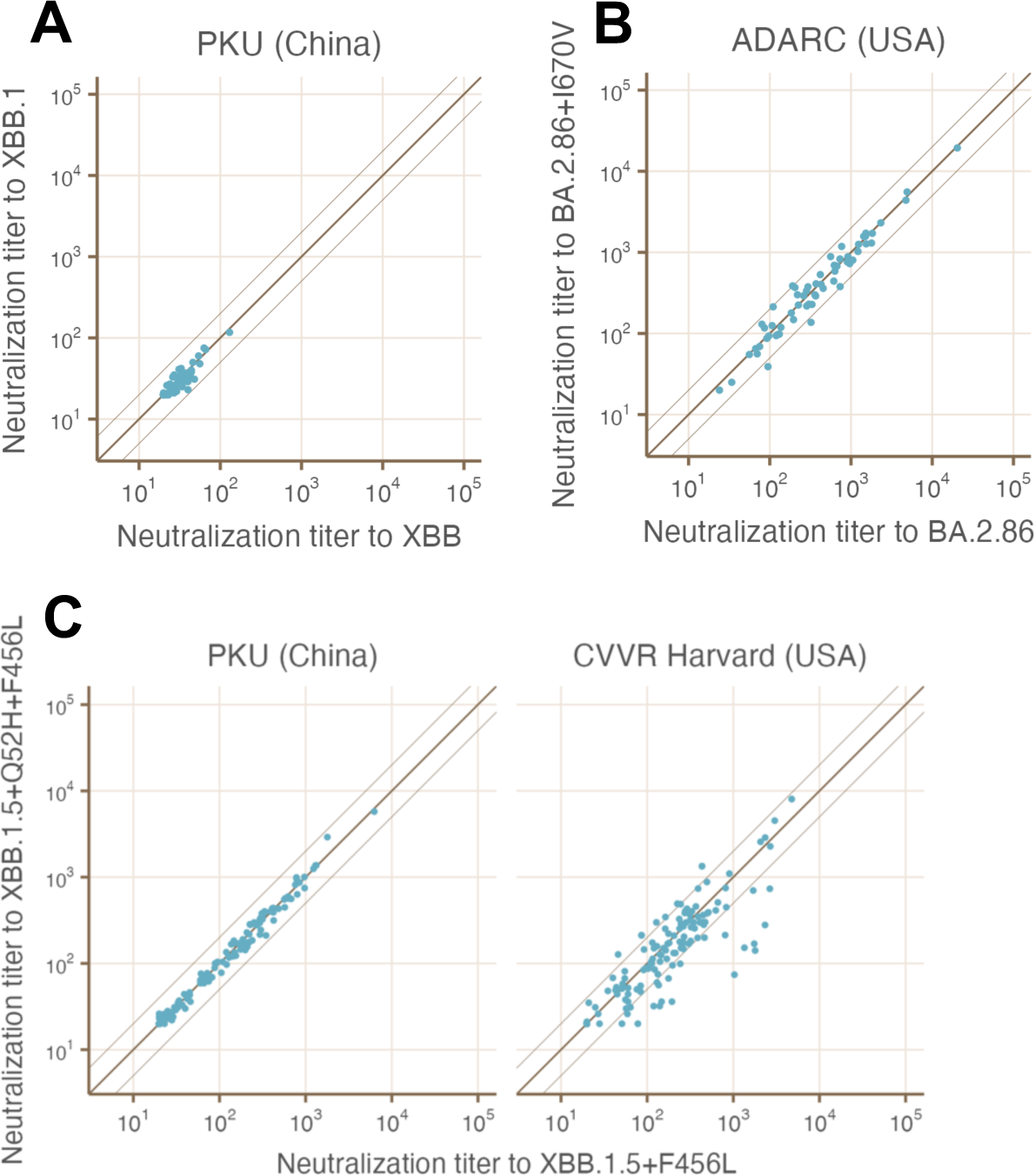
Comparison of titers to pairs of variants (**A:** BA.2.86 against BA.2.86+I670V; **B:** XBB against XBB.1; **C:** XBB.1.5+F456L against XBB.1.5+Q52H+F456L) which are combined into a single variant for later analysis, as indicated in “Grouping of variants” in Methods. Each serum which is titrated against both variants is plotted a single point. Faint diagonal lines indicate a 2-fold difference between the titers on the x and y axes. For each pair, the additional substitutions do not substantially affect neutralization, though a small number of sera from CVVR Harvard (USA) appear to be sensitive to the Q52H substitution in XBB.1.5+Q52H+F456L.

**Figure S6.**
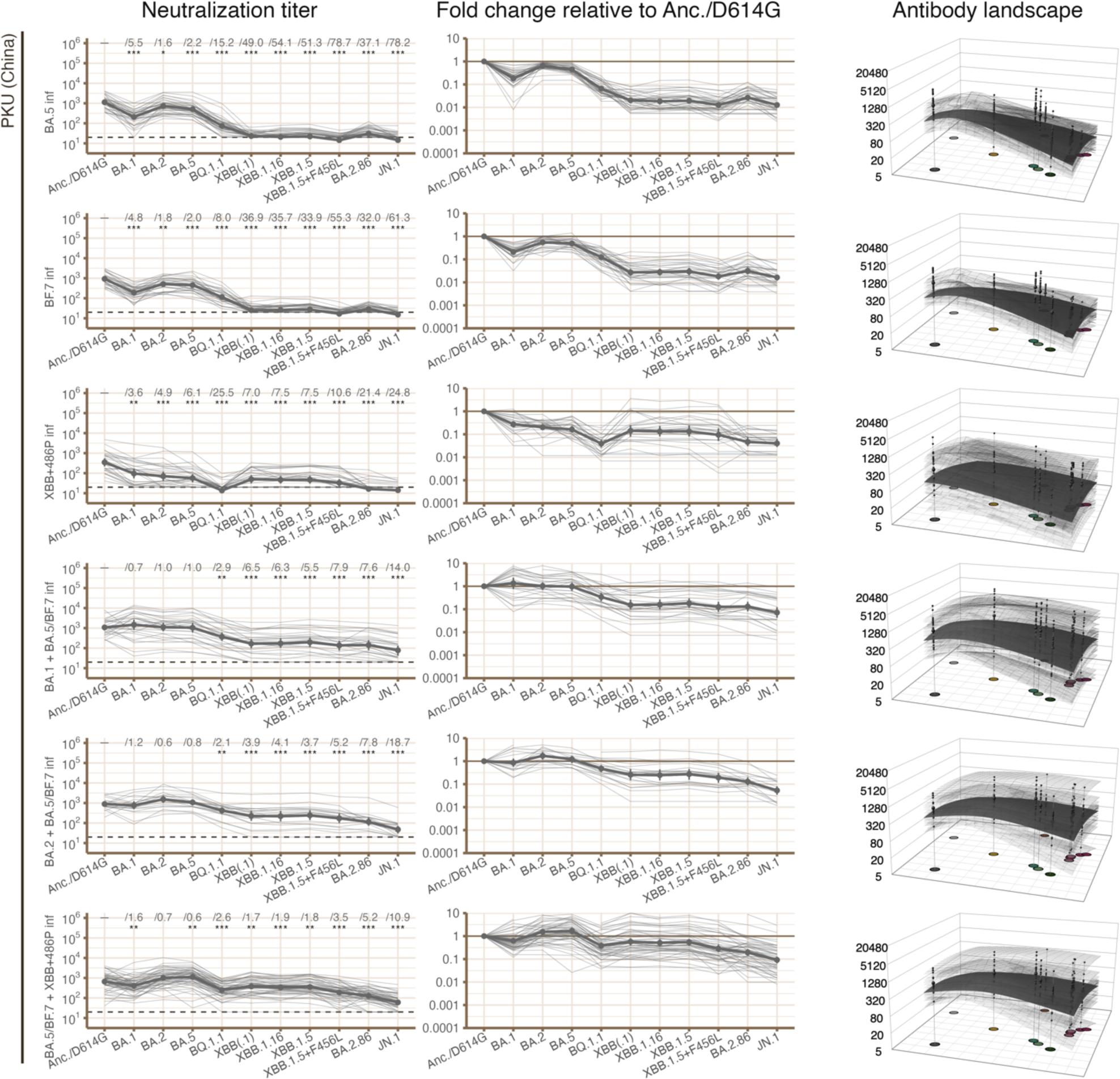
For each laboratory, neutralization titers (left column), fold change relative to the Anc./D614G variant (center column), and antibody landscapes (right column) are shown for individual sera in each serum group. For each serum group, bold lines indicate geometric mean titer and geometric mean fold change relative to the Anc./D614G variant, and the dark plane indicates the geometric mean antibody landscape. Pale lines and planes indicate these quantities for each individual serum within the serum group. In some serum groups from ADARC, titrations were not performed using BA.2.86, so titers were estimated from JN.1 titers for the purposes of producing antibody landscapes, as described in “Estimating BA.2.86 titers from JN.1 titers” in Methods. GMTs calculated from these estimated titers are included in Figure S9 as points connected with dashed lines. Fold change relative to Anc./D614G is written in the “Neutralization titer” column. Asterisks indicate level of statistical significance (* = p<0.05; ** = p<0.01; *** = p<0.001) as determined by the Wilcoxon paired difference test. 95% confidence intervals are shown, calculated by the non-parametric bootstrap using 1000 replicates. Abbreviations are used in serum group names and exposure history: “vax” = vaccinated; “inf” = infected; “Anc.” = ancestral/Wuhan-1 SARS-CoV-2 virus; “biv.” = bivalent.

**Figure S7.**
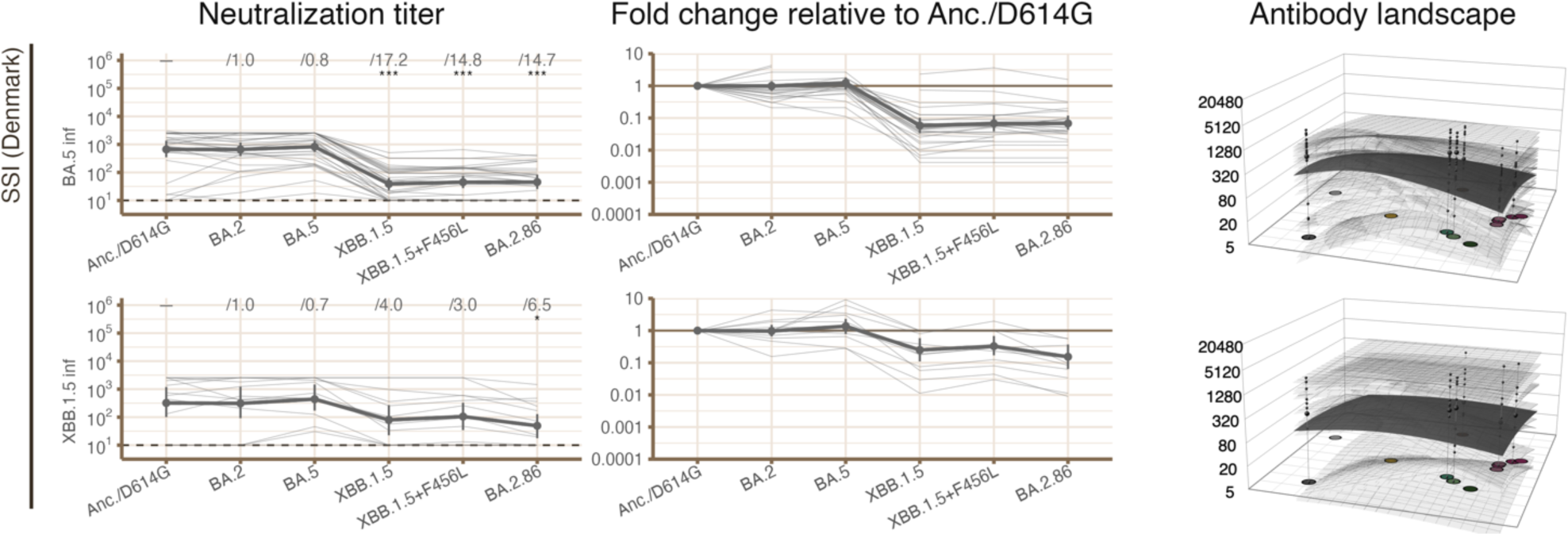
For each laboratory, neutralization titers (left column), fold change relative to the Anc./D614G variant (center column), and antibody landscapes (right column) are shown for individual sera in each serum group. For each serum group, bold lines indicate geometric mean titer and geometric mean fold change relative to the Anc./D614G variant, and the dark plane indicates the geometric mean antibody landscape. Pale lines and planes indicate these quantities for each individual serum within the serum group. In some serum groups from ADARC, titrations were not performed using BA.2.86, so titers were estimated from JN.1 titers for the purposes of producing antibody landscapes, as described in “Estimating BA.2.86 titers from JN.1 titers” in Methods. GMTs calculated from these estimated titers are included in Figure S9 as points connected with dashed lines. Fold change relative to Anc./D614G is written in the “Neutralization titer” column. Asterisks indicate level of statistical significance (* = p<0.05; ** = p<0.01; *** = p<0.001) as determined by the Wilcoxon paired difference test. 95% confidence intervals are shown, calculated by the non-parametric bootstrap using 1000 replicates. Abbreviations are used in serum group names and exposure history: “vax” = vaccinated; “inf” = infected; “Anc.” = ancestral/Wuhan-1 SARS-CoV-2 virus; “biv.” = bivalent.

**Figure S8.**
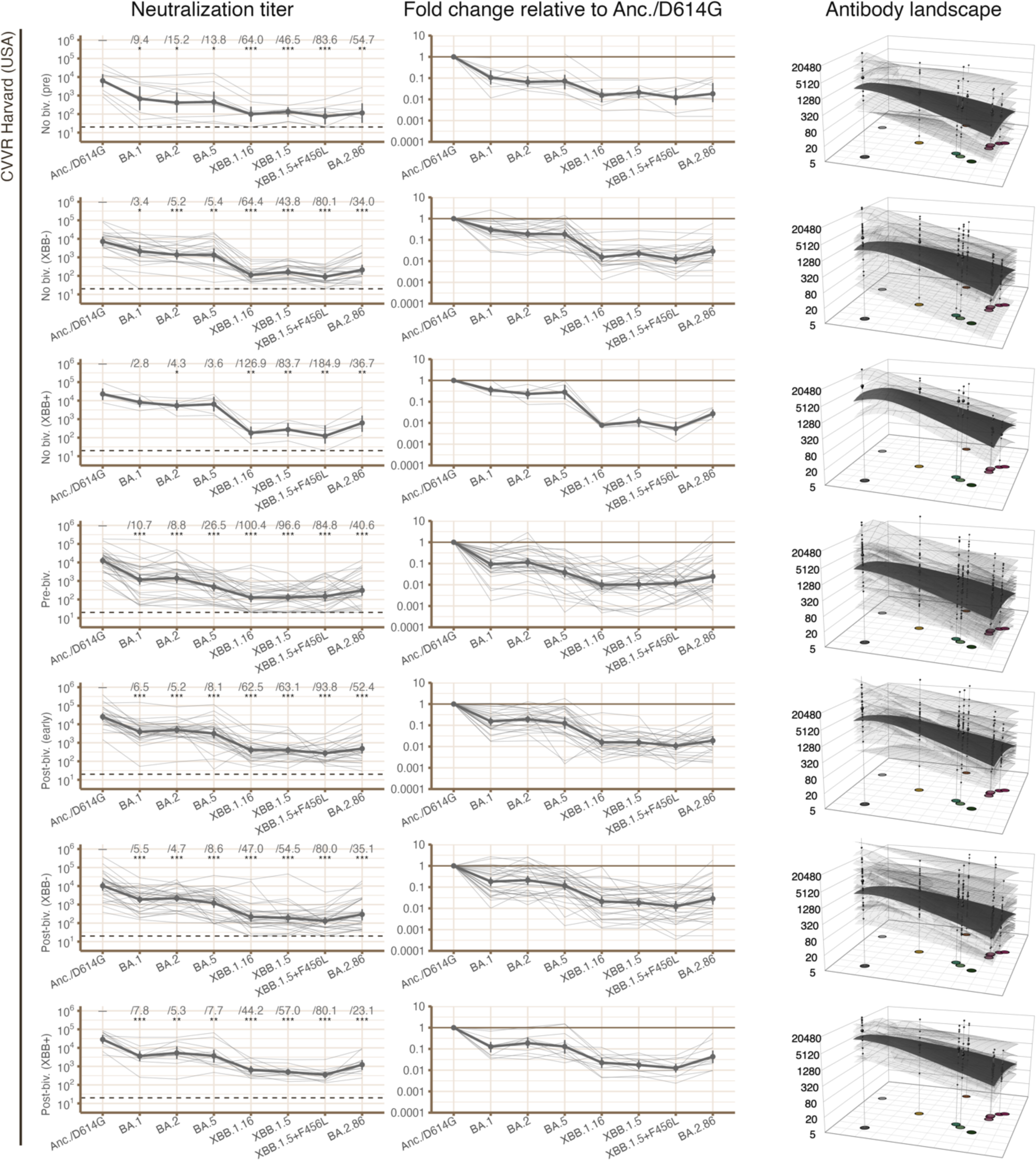
For each laboratory, neutralization titers (left column), fold change relative to the Anc./D614G variant (center column), and antibody landscapes (right column) are shown for individual sera in each serum group. For each serum group, bold lines indicate geometric mean titer and geometric mean fold change relative to the Anc./D614G variant, and the dark plane indicates the geometric mean antibody landscape. Pale lines and planes indicate these quantities for each individual serum within the serum group. In some serum groups from ADARC, titrations were not performed using BA.2.86, so titers were estimated from JN.1 titers for the purposes of producing antibody landscapes, as described in “Estimating BA.2.86 titers from JN.1 titers” in Methods. GMTs calculated from these estimated titers are included in Figure S9 as points connected with dashed lines. Fold change relative to Anc./D614G is written in the “Neutralization titer” column. Asterisks indicate level of statistical significance (* = p<0.05; ** = p<0.01; *** = p<0.001) as determined by the Wilcoxon paired difference test. 95% confidence intervals are shown, calculated by the non-parametric bootstrap using 1000 replicates. Abbreviations are used in serum group names and exposure history: “vax” = vaccinated; “inf” = infected; “Anc.” = ancestral/Wuhan-1 SARS-CoV-2 virus; “biv.” = bivalent.

**Figure S9.**
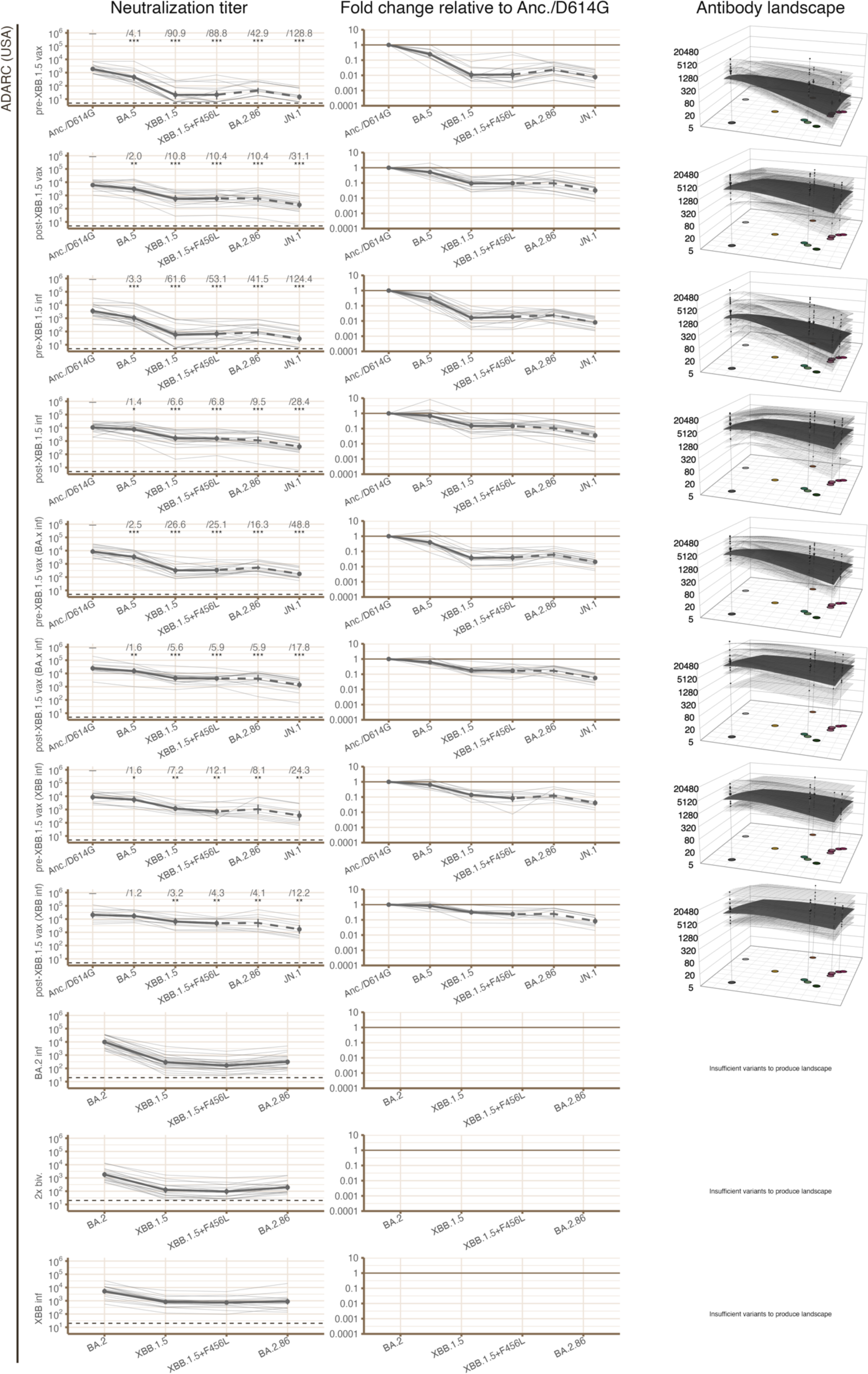
For each laboratory, neutralization titers (left column), fold change relative to the Anc./D614G variant (center column), and antibody landscapes (right column) are shown for individual sera in each serum group. For each serum group, bold lines indicate geometric mean titer and geometric mean fold change relative to the Anc./D614G variant, and the dark plane indicates the geometric mean antibody landscape. Pale lines and planes indicate these quantities for each individual serum within the serum group. In some serum groups from ADARC, titrations were not performed using BA.2.86, so titers were estimated from JN.1 titers for the purposes of producing antibody landscapes, as described in “Estimating BA.2.86 titers from JN.1 titers” in Methods. GMTs calculated from these estimated titers are included in Figure S9 as points connected with dashed lines. Fold change relative to Anc./D614G is written in the “Neutralization titer” column. Asterisks indicate level of statistical significance (* = p<0.05; ** = p<0.01; *** = p<0.001) as determined by the Wilcoxon paired difference test. 95% confidence intervals are shown, calculated by the non-parametric bootstrap using 1000 replicates. Abbreviations are used in serum group names and exposure history: “vax” = vaccinated; “inf” = infected; “Anc.” = ancestral/Wuhan-1 SARS-CoV-2 virus; “biv.” = bivalent.

**Figure S10.**
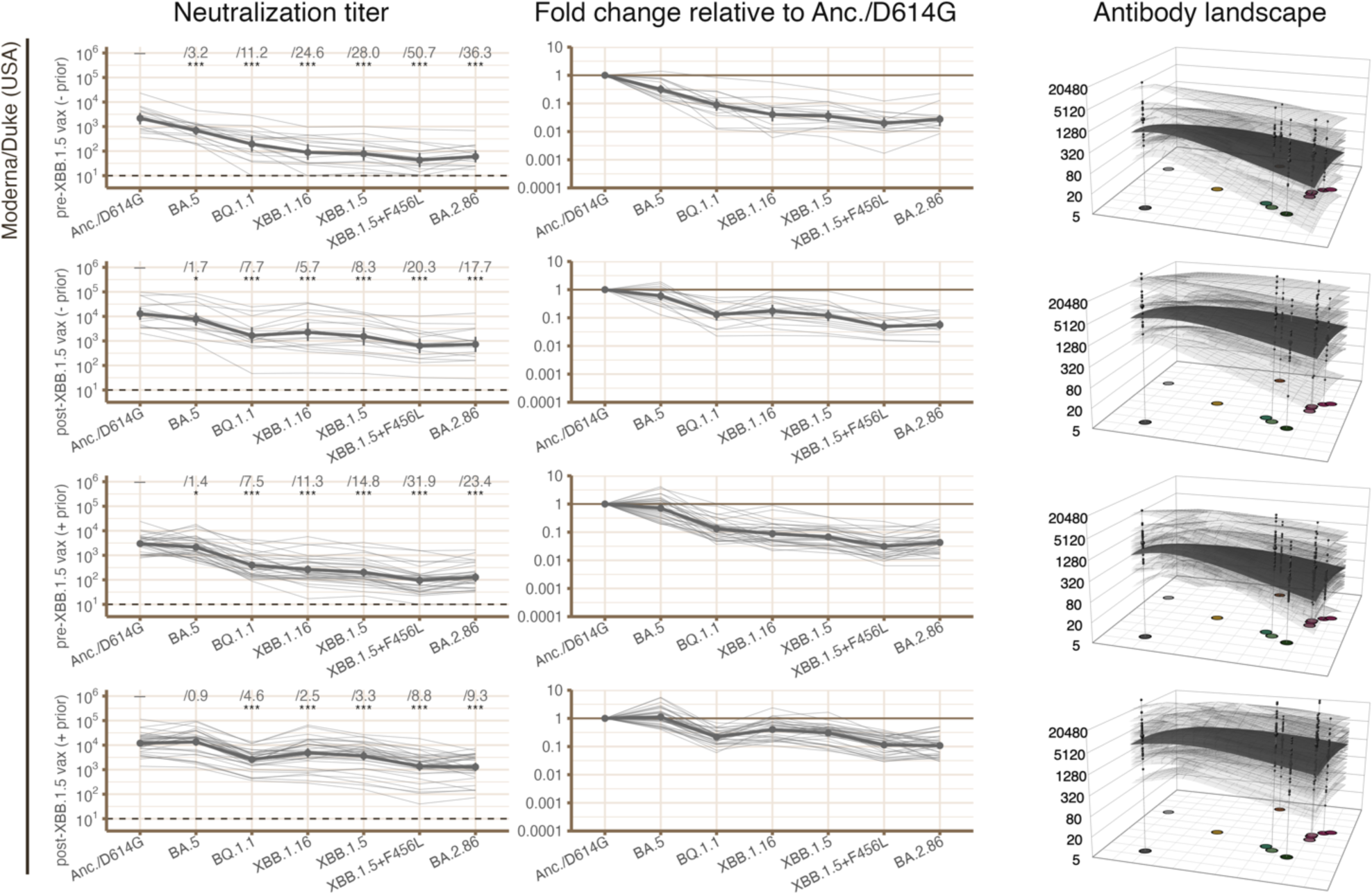
For each laboratory, neutralization titers (left column), fold change relative to the Anc./D614G variant (center column), and antibody landscapes (right column) are shown for individual sera in each serum group. For each serum group, bold lines indicate geometric mean titer and geometric mean fold change relative to the Anc./D614G variant, and the dark plane indicates the geometric mean antibody landscape. Pale lines and planes indicate these quantities for each individual serum within the serum group. In some serum groups from ADARC, titrations were not performed using BA.2.86, so titers were estimated from JN.1 titers for the purposes of producing antibody landscapes, as described in “Estimating BA.2.86 titers from JN.1 titers” in Methods. GMTs calculated from these estimated titers are included in Figure S9 as points connected with dashed lines. Fold change relative to Anc./D614G is written in the “Neutralization titer” column. Asterisks indicate level of statistical significance (* = p<0.05; ** = p<0.01; *** = p<0.001) as determined by the Wilcoxon paired difference test. 95% confidence intervals are shown, calculated by the non-parametric bootstrap using 1000 replicates. Abbreviations are used in serum group names and exposure history: “vax” = vaccinated; “inf” = infected; “Anc.” = ancestral/Wuhan-1 SARS-CoV-2 virus; “biv.” = bivalent.

**Figure S11.**
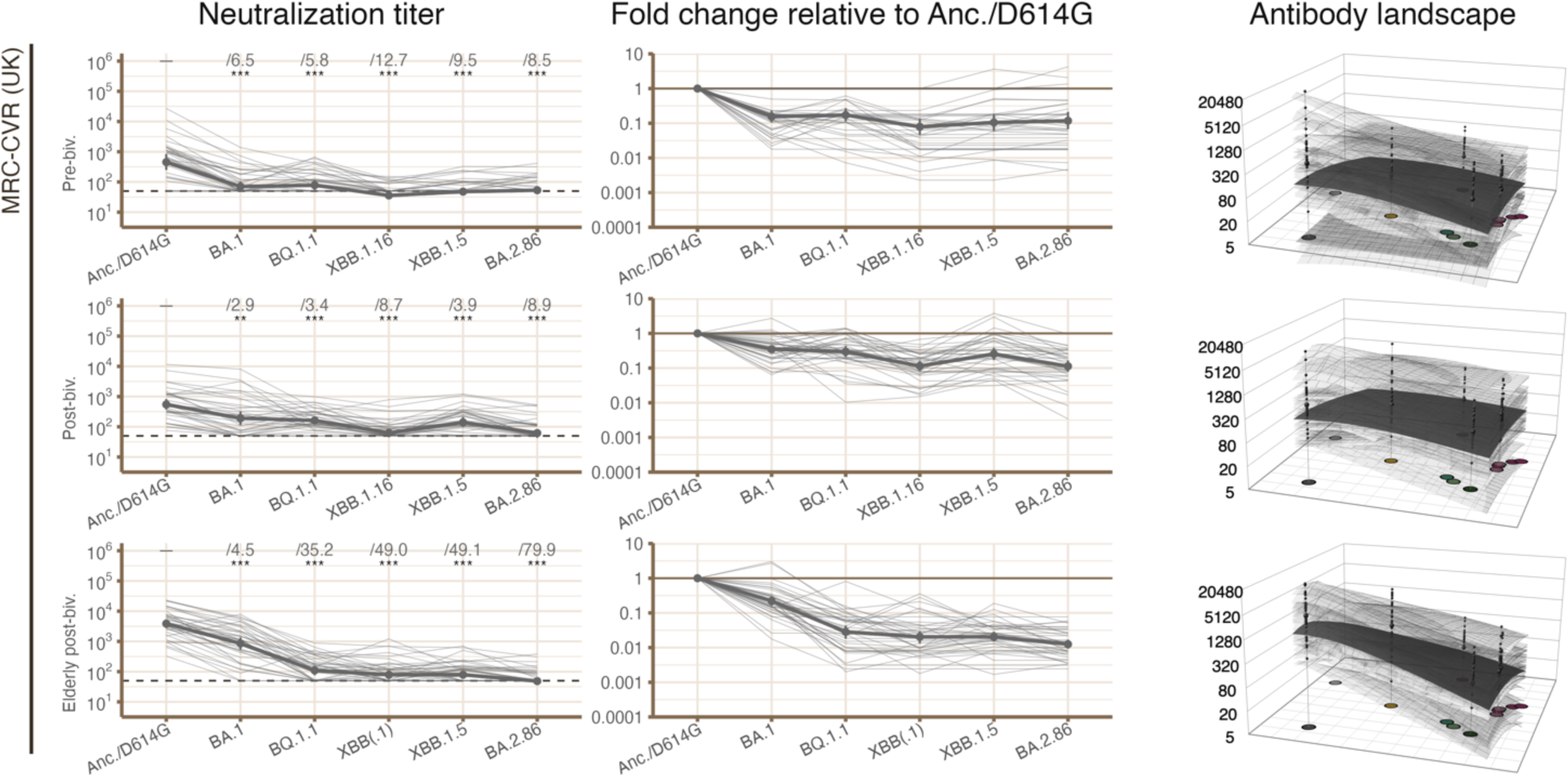
For each laboratory, neutralization titers (left column), fold change relative to the Anc./D614G variant (center column), and antibody landscapes (right column) are shown for individual sera in each serum group. For each serum group, bold lines indicate geometric mean titer and geometric mean fold change relative to the Anc./D614G variant, and the dark plane indicates the geometric mean antibody landscape. Pale lines and planes indicate these quantities for each individual serum within the serum group. In some serum groups from ADARC, titrations were not performed using BA.2.86, so titers were estimated from JN.1 titers for the purposes of producing antibody landscapes, as described in “Estimating BA.2.86 titers from JN.1 titers” in Methods. GMTs calculated from these estimated titers are included in Figure S9 as points connected with dashed lines. Fold change relative to Anc./D614G is written in the “Neutralization titer” column. Asterisks indicate level of statistical significance (* = p<0.05; ** = p<0.01; *** = p<0.001) as determined by the Wilcoxon paired difference test. 95% confidence intervals are shown, calculated by the non-parametric bootstrap using 1000 replicates. Abbreviations are used in serum group names and exposure history: “vax” = vaccinated; “inf” = infected; “Anc.” = ancestral/Wuhan-1 SARS-CoV-2 virus; “biv.” = bivalent.

**Figure S12.**
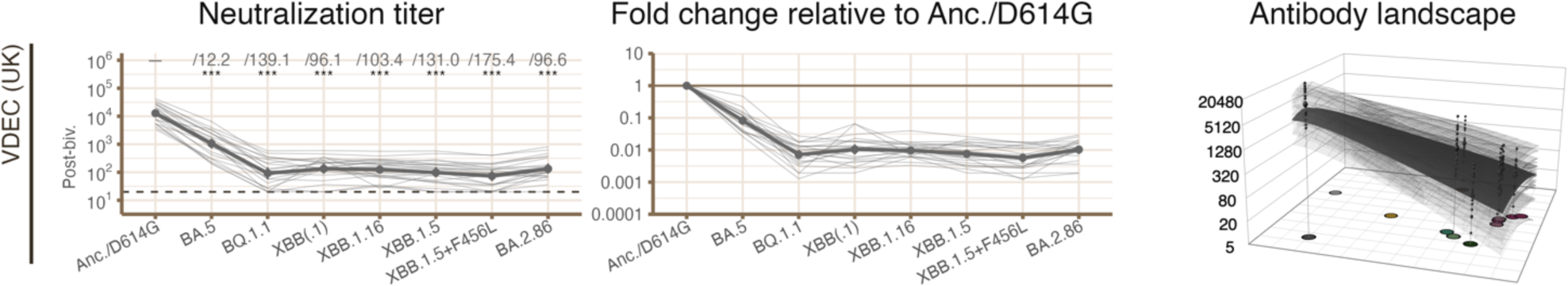
For each laboratory, neutralization titers (left column), fold change relative to the Anc./D614G variant (center column), and antibody landscapes (right column) are shown for individual sera in each serum group. For each serum group, bold lines indicate geometric mean titer and geometric mean fold change relative to the Anc./D614G variant, and the dark plane indicates the geometric mean antibody landscape. Pale lines and planes indicate these quantities for each individual serum within the serum group. In some serum groups from ADARC, titrations were not performed using BA.2.86, so titers were estimated from JN.1 titers for the purposes of producing antibody landscapes, as described in “Estimating BA.2.86 titers from JN.1 titers” in Methods. GMTs calculated from these estimated titers are included in Figure S9 as points connected with dashed lines. Fold change relative to Anc./D614G is written in the “Neutralization titer” column. Asterisks indicate level of statistical significance (* = p<0.05; ** = p<0.01; *** = p<0.001) as determined by the Wilcoxon paired difference test. 95% confidence intervals are shown, calculated by the non-parametric bootstrap using 1000 replicates. Abbreviations are used in serum group names and exposure history: “vax” = vaccinated; “inf” = infected; “Anc.” = ancestral/Wuhan-1 SARS-CoV-2 virus; “biv.” = bivalent.

**Figure S13.**
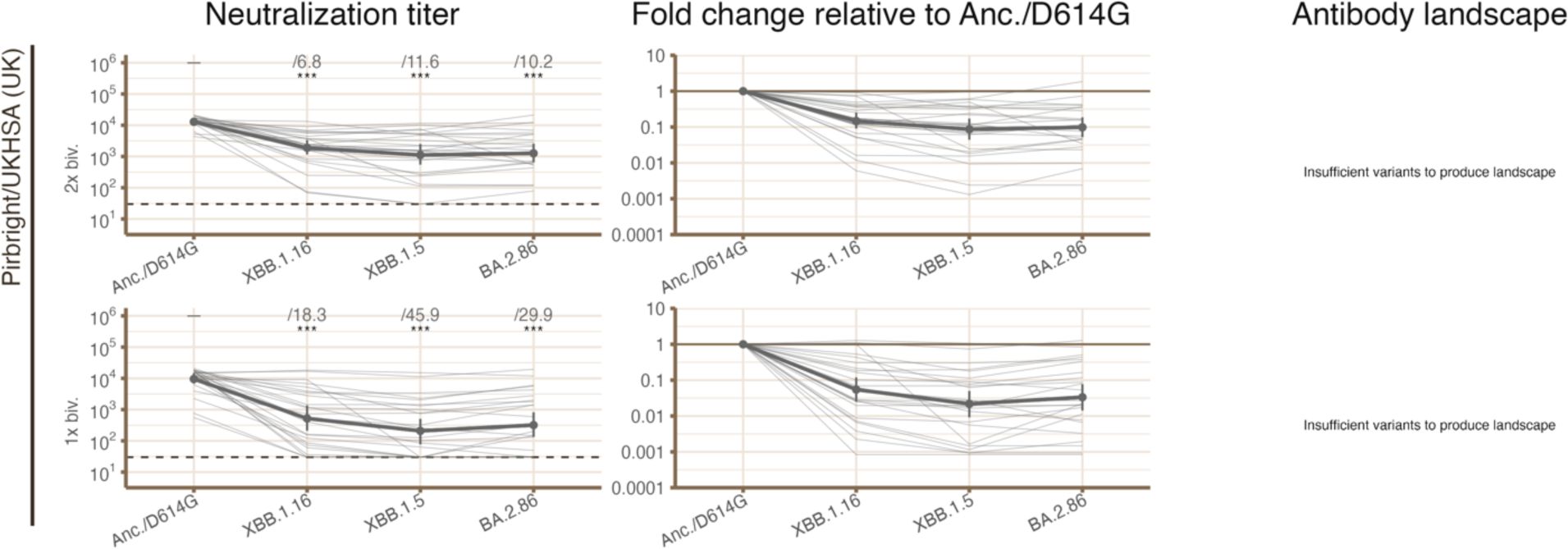
For each laboratory, neutralization titers (left column), fold change relative to the Anc./D614G variant (center column), and antibody landscapes (right column) are shown for individual sera in each serum group. For each serum group, bold lines indicate geometric mean titer and geometric mean fold change relative to the Anc./D614G variant, and the dark plane indicates the geometric mean antibody landscape. Pale lines and planes indicate these quantities for each individual serum within the serum group. In some serum groups from ADARC, titrations were not performed using BA.2.86, so titers were estimated from JN.1 titers for the purposes of producing antibody landscapes, as described in “Estimating BA.2.86 titers from JN.1 titers” in Methods. GMTs calculated from these estimated titers are included in Figure S9 as points connected with dashed lines. Fold change relative to Anc./D614G is written in the “Neutralization titer” column. Asterisks indicate level of statistical significance (* = p<0.05; ** = p<0.01; *** = p<0.001) as determined by the Wilcoxon paired difference test. 95% confidence intervals are shown, calculated by the non-parametric bootstrap using 1000 replicates. Abbreviations are used in serum group names and exposure history: “vax” = vaccinated; “inf” = infected; “Anc.” = ancestral/Wuhan-1 SARS-CoV-2 virus; “biv.” = bivalent.

**Table S1:**

| Laboratory | Pseudovirus backbone or live virus | Target cell type | Full length or truncated spike |
| --- | --- | --- | --- |
| PKU | VSV pseudovirus | Huh-7 | Full length spike |
| SSI | Live virus | Vero-E6 | N/A |
| MRC-CVR | Lentivirus pseudovirus | HEK293-ACE2 | Full length spike |
| VDEC | Live virus | Vero-E6 | N/A |
| Pirbright/UKHSA | Lentivirus pseudovirus | HEK293T-ACE2 | CT-truncated spike |
| CVVR Harvard | Lentivirus pseudovirus | HEK293T-ACE2 | CT-truncated spike |
| ADARC | VSV pseudovirus | Vero-E6 | CT-truncated spike |
| Moderna/Duke | Lentivirus pseudovirus | HEK293T-ACE2 | Full length spike |

Additional summary information about assay protocols used in each laboratory. Full assay protocols are given in the “Individual group methodologies” section of Methods. “CT-truncated spike” refers to the removal of the cytoplasmic tail of the spike protein to increase the spike density and infectivity of pseudoviruses (Yu et al. 2021).

## References

Bewley, Kevin R., Naomi S. Coombes, Luc Gagnon, Lorna McInroy, Natalie Baker, Imam Shaik, Julien R. St-Jean, et al. 2021. “Quantification of SARS-CoV-2 Neutralizing Antibody by Wild-Type Plaque Reduction Neutralization, Microneutralization and Pseudotyped Virus Neutralization Assays.” Nature Protocols 16 (6): 3114–40.

Chalkias, Spyros, Frank Eder, Brandon Essink, Shishir Khetan, Biliana Nestorova, Jing Feng, Xing Chen, et al. 2022. “Safety, Immunogenicity and Antibody Persistence of a Bivalent Beta-Containing Booster Vaccine against COVID-19: A Phase 2/3 Trial.” Nature Medicine 28 (11): 2388–97.

Chalkias, Spyros, Nichole McGhee, Jordan L. Whatley, Brandon Essink, Adam Brosz, Joanne E. Tomassini, Bethany Girard, et al. 2023. “Safety and Immunogenicity of XBB.1.5-Containing MRNA Vaccines.” BioRxiv. 10.1101/2023.08.22.23293434.

Chen, Chaoran, Sarah Nadeau, Michael Yared, Philippe Voinov, Ning Xie, Cornelius Roemer, and Tanja Stadler. 2022. “CoV-Spectrum: Analysis of Globally Shared SARS-CoV-2 Data to Identify and Characterize New Variants.” Bioinformatics 38 (6): 1735–37.

Coombes, Naomi S., Kevin R. Bewley, Yann Le Duff, Nassim Alami-Rahmouni, Kathryn A. Ryan, Sarah Kempster, Deborah Ferguson, et al. 2023. “Evaluation of the Neutralising Antibody Response in Human and Hamster Sera against SARS-CoV-2 Variants up to and Including BA.2.86 Using an Authentic Virus Neutralisation Assay.” BioRxiv. 10.1101/2023.10.21.563398.

Coombes, Naomi S., Kevin R. Bewley, Yann Le Duff, Matthew Hurley, Lauren J. Smith, Thomas M. Weldon, Karen Osman, et al. 2023. “Assessment of the Biological Impact of SARS-CoV-2 Genetic Variation Using an Authentic Virus Neutralisation Assay with Convalescent Plasma, Vaccinee Sera, and Standard Reagents.” Viruses 15 (3). 10.3390/v15030633.

Edouard Mathieu, Hannah Ritchie, Lucas Rodés-Guirao, Cameron Appel, Charlie Giattino, Joe Hasell, Bobbie Macdonald, Saloni Dattani, Diana Beltekian, Esteban Ortiz-Ospina and Max Roser. n.d. “Coronavirus Pandemic (COVID-19).” Ourworldindata.org. Accessed August 1, 2024. https://ourworldindata.org/coronavirus.

Frische, Anders, Patrick Terrence Brooks, Mikkel Gybel-Brask, Susanne Gjørup Sækmose, Bitten Aagaard Jensen, Susan Mikkelsen, Mie Topholm Bruun, et al. 2022. “Optimization and Evaluation of a Live Virus SARS-CoV-2 Neutralization Assay.” PloS One 17 (7): e0272298.

Gilbert, Peter B., David C. Montefiori, Adrian B. McDermott, Youyi Fong, David Benkeser, Weiping Deng, Honghong Zhou, et al. 2022. “Immune Correlates Analysis of the MRNA-1273 COVID-19 Vaccine Efficacy Clinical Trial.” *Science (New York*, N.Y.) 375 (6576): 43–50.

Jacobsen, Henning, Ioannis Sitaras, Maeva Katzmarzyk, Viviana Cobos Jiménez, Robert Naughton, Melissa M. Higdon, and Maria Deloria Knoll. 2023. “Systematic Review and Meta-Analysis of the Factors Affecting Waning of Post-Vaccination Neutralizing Antibody Responses against SARS-CoV-2.” Npj Vaccines 8 (1): 1–6.

Kaku, Yu, Kaho Okumura, Miguel Padilla-Blanco, Yusuke Kosugi, Keiya Uriu, Alfredo A. Hinay Jr, Luo Chen, et al. 2024. “Virological Characteristics of the SARS-CoV-2 JN.1 Variant.” The Lancet Infectious Diseases 24 (2): e82.

Khan, Khadija, Gila Lustig, Cornelius Römer, Kajal Reedoy, Zesuliwe Jule, Farina Karim, Yashica Ganga, et al. 2023. “Evolution and Neutralization Escape of the SARS-CoV-2 BA.2.86 Subvariant.” Nature Communications 14 (1): 1–9.

Khare, Shruti, GISAID Global Data Science Initiative (GISAID), Munich, Germany, Céline Gurry, Lucas Freitas, Mark B Schultz, Gunter Bach, Amadou Diallo, et al. 2021. “GISAID’s Role in Pandemic Response.” China CDC Weekly 3 (49): 1049–51.

Lasrado, Ninaad, Ai-Ris Y. Collier, Nicole P. Hachmann, Jessica Miller, Marjorie Rowe, Eleanor D. Schonberg, Stefanie L. Rodrigues, et al. 2023. “Neutralization Escape by SARS-CoV-2 Omicron Subvariant BA.2.86.” Vaccine 41 (47): 6904–9.

Lasrado, Ninaad, Ai-Ris Y. Collier, Jessica Miller, Nicole P. Hachmann, Jinyan Liu, Michaela Sciacca, Cindy Wu, et al. 2023. “Waning Immunity against XBB.1.5 Following Bivalent MRNA Boosters.” *BioRxiv.Org: The Preprint Server for Biology*, January. 10.1101/2023.01.22.525079.

Lassaunière, Ria, Charlotta Polacek, Magdalena Utko, Karina M. Sørensen, Sharmin Baig, Kirsten Ellegaard, Leandro A. Escobar-Herrera, et al. 2023. “Virus Isolation and Neutralisation of SARS-CoV-2 Variants BA.2.86 and EG.5.1.” The Lancet Infectious Diseases 23 (12): e509–10.

Newman, Joseph, Nazia Thakur, Thomas P. Peacock, Dagmara Bialy, Ahmed M. E. Elrefaey, Carlijn Bogaardt, Daniel L. Horton, et al. 2022. “Neutralizing Antibody Activity against 21 SARS-CoV-2 Variants in Older Adults Vaccinated with BNT162b2.” Nature Microbiology 7 (8): 1180–88.

R Core Team. 2022. “R: A Language and Environment for Statistical Computing.” Vienna, Austria: R Foundation for Statistical Computing. https://www.R-project.org/.

Rössler, Annika, Antonia Netzl, Ludwig Knabl, David Bante, Samuel H. Wilks, Wegene Borena, Dorothee von Laer, Derek J. Smith, and Janine Kimpel. 2023. “Characterizing SARS-CoV-2 Neutralization Profiles after Bivalent Boosting Using Antigenic Cartography.” Nature Communications 14 (1): 5224.

Ryan, Kathryn A., Kevin R. Bewley, Robert J. Watson, Christopher Burton, Oliver Carnell, Breeze E. Cavell, Amy Challis, et al. 2023. “Syrian Hamster Convalescence from Prototype SARS-CoV-2 Confers Measurable Protection against the Attenuated Disease Caused by the Omicron Variant.” PLoS Pathogens 19 (4): e1011293.

Shu, Yuelong, and John McCauley. 2017. “GISAID: Global Initiative on Sharing All Influenza Data – from Vision to Reality.” Euro Surveillance: Bulletin Europeen Sur Les Maladies Transmissibles = European Communicable Disease Bulletin 22 (13). 10.2807/1560-7917.es.2017.22.13.30494.

Smith, Derek J., Alan S. Lapedes, Jan C. de Jong, Theo M. Bestebroer, Guus F. Rimmelzwaan, Albert D. M. E. Osterhaus, and Ron A. M. Fouchier. 2004. “Mapping the Antigenic and Genetic Evolution of Influenza Virus.” Science 305 (5682): 371–76.

Wang, Qian, Yicheng Guo, Anthony Bowen, Ian A. Mellis, Riccardo Valdez, Carmen Gherasim, Aubree Gordon, Lihong Liu, and David D. Ho. 2023. “XBB.1.5 Monovalent MRNA Vaccine Booster Elicits Robust Neutralizing Antibodies against Emerging SARS-CoV-2 Variants.” BioRxiv. 10.1101/2023.11.26.568730.

Wang, Qian, Yicheng Guo, Liyuan Liu, Logan T. Schwanz, Zhiteng Li, Manoj S. Nair, Jerren Ho, et al. 2023. “Antigenicity and Receptor Affinity of SARS-CoV-2 BA.2.86 Spike.” *Nature*, October. 10.1038/s41586-023-06750-w.

Wilks, Sam. 2023. “Racmacs: Antigenic Cartography Macros.” https://acorg.github.io/Racmacs/.

Wilks, Samuel. 2021. “Ablandscapes: Making Antibody Landscapes Using R.” https://github.com/acorg/ablandscapes.

Willett, Brian J., Joe Grove, Oscar A. MacLean, Craig Wilkie, Giuditta De Lorenzo, Wilhelm Furnon, Diego Cantoni, et al. 2022. “SARS-CoV-2 Omicron Is an Immune Escape Variant with an Altered Cell Entry Pathway.” Nature Microbiology 7 (8): 1161–79.

Willett, Brian J., Nicola Logan, Sam Scott, Chris Davis, Therese McSorley, Patawee Asamaphan, Margaret J. Hosie, et al. 2023. “Omicron BA.2.86 Cross-Neutralising Activity in Community Sera from the UK.” The Lancet 402 (10417): 2075–76.

World Health Organisation. 2023. “Initial Risk Evaluation of BA.2.86 and Its Sublineages.”

World Health Organization. n.d. “WHO Coronavirus (COVID-19) Dashboard.” Data.who.int. Accessed August 1, 2024. https://data.who.int/dashboards/covid19/about.

Yang, Sijie, Yuanling Yu, Fanchong Jian, Weiliang Song, Ayijiang Yisimayi, Xiaosu Chen, Yanli Xu, et al. 2023. “Antigenicity and Infectivity Characterisation of SARS-CoV-2 BA.2.86.” The Lancet Infectious Diseases 23 (11): e457–59.

Yang, Sijie, Yuanling Yu, Yanli Xu, Fanchong Jian, Weiliang Song, Ayijiang Yisimayi, Peng Wang, et al. 2024. “Fast Evolution of SARS-CoV-2 BA.2.86 to JN.1 under Heavy Immune Pressure.” The Lancet Infectious Diseases 24 (2): e70–72.

Yisimayi, Ayijiang, Weiliang Song, Jing Wang, Fanchong Jian, Yuanling Yu, Xiaosu Chen, Yanli Xu, et al. 2024. “Repeated Omicron Exposures Override Ancestral SARS-CoV-2 Immune Imprinting.” Nature 625 (7993): 148–56.

Yu, Jingyou, Zhenfeng Li, Xuan He, Makda S. Gebre, Esther A. Bondzie, Huahua Wan, Catherine Jacob-Dolan, et al. 2021. “Deletion of the SARS-CoV-2 Spike Cytoplasmic Tail Increases Infectivity in Pseudovirus Neutralization Assays.” Journal of Virology 95 (11). 10.1128/jvi.00044-21.

